# Airway organoids reveal patterns of Influenza A tropism and adaptation in wildlife species

**DOI:** 10.64898/2025.12.17.694819

**Authors:** Ferran Tarrés-Freixas, Gerardo Ceada, Francesc Català-Moll, Maria Casadellà, Sara Tolosa-Alarcón, Eloi Franco-Trepat, Rytis Boreika, Vanessa Almagro-Delgado, Noelia Carmona, Josep Estruch, Hugo Fernández-Bellon, Laura Fuentes-Moyano, Rafael Molina-López, Nuria Navarro, Mariona Parera, Marta Pérez-Simó, Joan Repullés, Núria Roca, Odalys Torné, Roser Velarde, Alfonso Valencia, Roger Paredes, Nuria Izquierdo-Useros, Miguel Romero-Durana, Karl Kochanowski, Júlia Vergara-Alert

**Affiliations:** Unitat Mixta d’investigació IRTA-UAB en Sanitat Animal, CReSA, UAB, Bellaterra, 08193, Spain; IRTA. Programa de Sanitat Animal. Centre de Recerca en Sanitat Animal (CReSA). Campus de la Universitat Autònoma de Barcelona (UAB), Bellaterra, 08193, Spain; WOAH (OIE) Collaborating Centre for the Research and Control of Emerging and Re-emerging Swine Diseases in Europe (IRTA-CReSA), Bellaterra, 08193, Spain; Biosciences Department, Faculty of Sciences and Technology, University of Vic - Central University of Catalonia, Vic, 08500, Spain; Department of Cell Biology, Physiology and Immunology, Universitat Autònoma de Barcelona, Bellaterra, 08193, Spain; IrsiCaixa,. Germans Trias i Pujol Research Institute (IGTP), Universitat Autònoma de Barcelona (UAB) Badalona, 08916, Spain; CIBER of Precision Medicine Against Antimicrobial Resistance (MePRAM), Carlos III Health Institute, Madrid, 28029, Spain; Computational Biology Group, Life Sciences Department, Barcelona Supercomputing Center (BSC), Barcelona, 08034, Spain; Zoo Barcelona, Parc Ciutadella s/n, Barcelona, 08003, Spain; Servei d’Ecopatologia de Fauna Salvatge (SEFaS) and Wildlife Ecology and Health group (WE&H), Departament de Medicina i Cirurgia Animals, Facultat de Veterinària, Campus de la Universitat Autònoma de Barcelona (UAB), Bellaterra, 08193, Spain; Centre de Fauna de Torreferrussa, Àrea de Gestió Ambiental Servei de Fauna i Flora, Forestal Catalana, 08130, Santa Perpètua de Mogoda, Barcelona, Spain; Institució Catalana de Recerca i Estudis Avançats (ICREA), Barcelona, 08010, Spain; CIBER Enfermedades Infecciosas (CIBERINFEC), Instituto de Salud Carlos III, Madrid, 28029, Spain; Germans Trias i Pujol Research Institute (IGTP), Universitat Autònoma de Barcelona (UAB) Badalona, 08916, Spain

**Keywords:** Influenza virus, Airway Organoids, Wildlife species, host tropism, H5N1 HPAIV, H1N1(pdm09)

## Abstract

Identifying animal species that are susceptible to the plethora of existing and emerging viruses is critical for predicting and containing disease outbreaks. Current efforts to assess viral tropism largely rely on experimental infection models, but such experiments are logistically and ethically infeasible for many wildlife species. To tackle this challenge, we developed a panel of airway organoids from ten taxonomically diverse wildlife and livestock species and evaluated their susceptibility to influenza viruses of mammalian (pH1N1) and avian (H5N1) origin. Our analyses revealed large species-specific differences in infection rate and cytopathogenicity that aligned with known *in vivo* data and field observations. Furthermore, we demonstrated that this organoid panel can serve as a powerful tool to elucidate receptor-binding mechanisms, viral dynamics, and early host adaptation in poorly characterized animal species. In summary, this work provides a robust and ethically viable approach for evaluating viral tropism and adaptation in wildlife species, and fills a critical gap in current pandemic preparedness, zoonotic disease surveillance, and wildlife conservation efforts.

## Introduction

Emerging and re-emerging infectious diseases represent a persistent threat to both humans and animals and impose substantial health, ecological, and economic burdens^1,2^. The increasing interconnection of human and animal populations, exacerbated by globalization and climate change, raises the chances of zoonotic spillover and limits outbreak control^3^. Therefore, there is a critical need to identify both susceptible populations and potential animal reservoirs for infectious agents^4^.

Traditionally, experimental infections in animals are considered the gold standard for evaluating host susceptibility, providing direct insights into pathogenicity and transmission dynamics^5,6^ However, this approach is unfeasible for many wildlife species, particularly for endangered ones, due to ethical constraints, logistical and technical challenges, and conservation efforts. These limitations have motivated the development of alternative *in vitro* approaches, such as the use of species-specific cell cultures^7–10^ or engineered cells expressing host receptors from different animals^11^, which enable controlled studies of viral dynamics under laboratory conditions. However, the availability of such *in vitro* models remains limited for many wildlife species. Moreover, many existing culture models –which often rely on immortalized cell lines– lack the cellular heterogeneity and interactions found in tissues, leading to discrepancies between *in vitro* findings and *in vivo* outcomes^12–16^.

In this work, we aimed to explore the use of organoids as an alternative approach for studying wildlife species’ susceptibility to infectious pathogens. These self-organizing, tissue-derived systems mimic the multicellular organization and function of their tissue of origin, thus bridging the gap between conventional cell cultures and *in vivo* models^17^. While organoids have proven their worth in human and murine models^18,19^, and are increasingly being used to study infectious diseases in livestock^20–24^, their application to wildlife remains largely unexplored. Recent efforts, such as bat-derived organoids to explore zoonotic reservoirs^25,26^, highlight the potential of organoids for studying viral infections in closely related wildlife species. However, whether such approaches also serve to directly compare viral tropism in more distantly related wildlife species is currently unclear, and efforts towards this end are hampered by difficulties in tissue access and protocol standardization.

To overcome these challenges, we developed a standardized procedure for generating and characterizing respiratory organoids from wildlife specimens. Using this approach, we successfully established airway organoids from ten taxonomically diverse wildlife and livestock species. As a case study, we infected these organoids with two influenza A virus strains: the 2009 pandemic H1N1 (pH1N1) and a clade 2.3.4.4b H5N1 high-pathogenicity avian influenza virus (HPAIV) strain. These infections revealed large differences in susceptibility to both viruses across species, especially when assessing infection rate and cytopathogenicity, which parallelled existing *in vivo* data and field observations. Nevertheless, we found that all tested organoid cultures were able to produce infective viral particles, and sequence analysis identified several instances of species-specific viral adaptation. Overall, this study offers a valuable and ethically viable approach for assessing host susceptibility to viral infection and viral adaptation in wildlife species, and contributes to broader efforts in pandemic preparedness, wildlife conservation, and zoonotic disease research.

## Results

### Section 1: Generation and morphological characterization of airway organoids from wildlife and livestock species

We first assessed whether a single and standardized protocol could reliably generate airway organoids from tissue samples obtained from taxonomically diverse animal species. We collected airway tissue samples from adult animals that were euthanized for welfare reasons. All samples were sourced from facilities in Catalonia housing both exotic and endemic wildlife species. We further included two livestock species (domestic pig and chicken) with well-characterized susceptibility patterns to various influenza A virus strains^27–30^ as reference. In total, our organoid panel encompassed ten species (8 mammalian and 2 avian), including endangered ones. Notably, for many of these species, organoid models (airway or otherwise) had not been previously reported (**Table 1**).

**Table 1.**
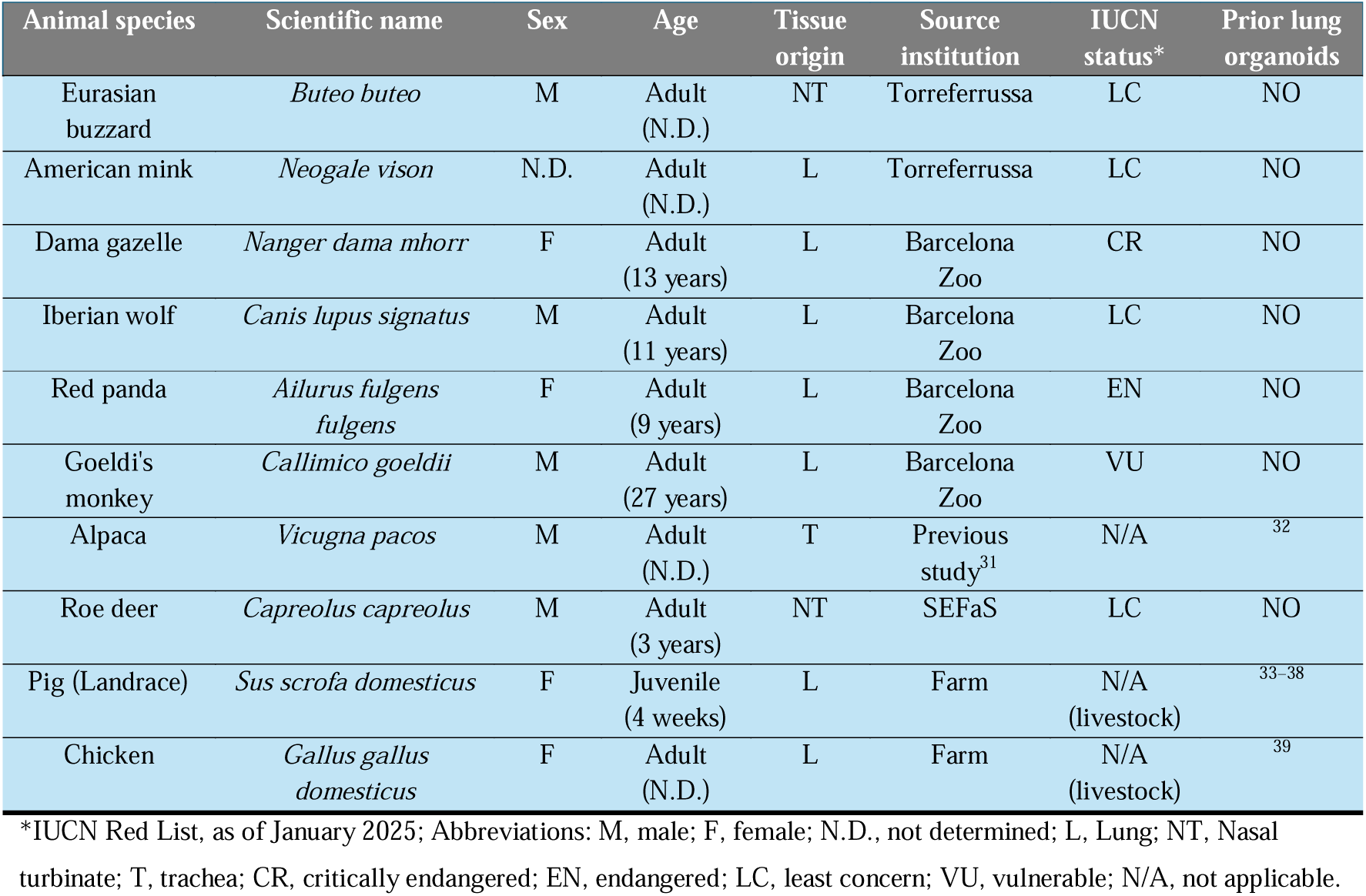
Overview of animal species used for airway organoid generation. The table includes both wildlife and livestock species included in the study, detailing sample origin, sex, tissue type, and conservation status (IUCN Red List, January 2025).

To generate organoids from these airway tissue samples, we followed a generalized protocol originally optimized for human tissue^40,41^, involving tissue dissociation, cell isolation, embedding in extracellular matrix (ECM), and subsequent organoid expansion in 3D culture (**Figure 1A**). Using this approach, we successfully established airway organoids from all ten species. All species yielded viable organoids within a few weeks (**Figure 1B**), which could be expanded over multiple passages. Brightfield microscopy revealed that organoids from all species exhibited characteristic spherical and cystic structures (**Figure 1B**). Although organoid size and density varied between species, the observed morphology was consistent with apical-in airway-derived epithelial structures^40,41^.

**Figure 1.**
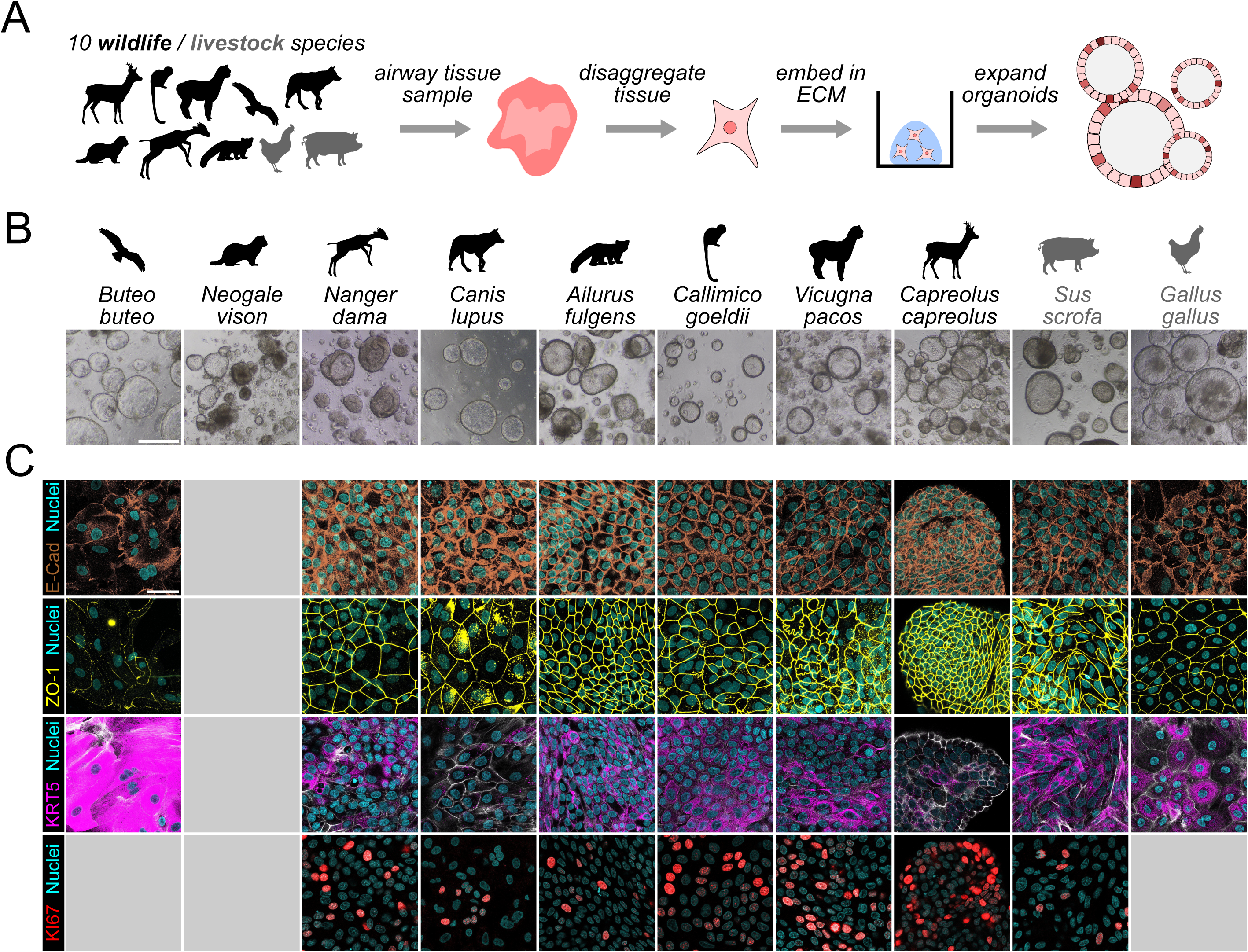
Establishing a panel of airway organoids from diverse wildlife and livestock species. **A)** Schematic approach. **B)** Example brightfield microscopy images of 3D airway organoids derived from each animal species. Silhouettes represent the corresponding species, with wildlife species in black and livestock species in gray. Scale bar: 400 µm (same scale in all images). **C)** Example immunofluorescence images of airway epithelial tissue markers in organoid monolayers. Gray boxes: Missing data due to technical reasons. Gray staining in KRT5: phalloidin. Scale bar: 40 µm (same scale in all images).

The apical-in polarity of 3D organoid cultures represents a technical challenge in organoid-based infection models. That is, the apical side of the cells - which is the primary entry site for respiratory viruses such as influenza viruses – faces the inaccessible internal lumen (see example in **Supplementary video 1**), and current approaches to make this apical side available, such as organoid disruption ^14^ or apical-out organoids^42,43^, are not yet optimized for wildlife-derived organoids. To circumvent this issue, we followed previous efforts and generated 2D monolayers from each 3D organoid culture in a 96-well plate format that ensure direct access to the apical surface^34,44–46^.

To assess whether these organoid-derived 2D monolayers have the expected molecular features of airway epithelial tissues, we performed immunofluorescence staining of key epithelial markers^34,47^ (**Figure 1C**). Across all tested species, we observed robust expression of E-cadherin (epithelial cell-cell adhesion marker, E-Cad), ZO-1 (tight junctions), Keratin 5 (basal cells, KRT5), and Ki67-positive nuclei (proliferative cells). Moreover, live imaging of the 2D-monolayers revealed the active cilia movement of ciliated cells (see example in **Supplementary video 2**). Thus, these findings confirm that the organoid-derived 2D monolayers we established from various animal species do retain key cell types and molecular features of the native airway epithelium.

### Section 2: Host-dependent differences in susceptibility to pH1N1 and H5N1

To investigate species-specific susceptibility to influenza infection, we exposed airway organoid monolayers derived from diverse animal species to two Influenza A viruses: the H1N1 strain responsible for the 2009 pandemics (mammalian origin, from now on **pH1N1**), and a H5N1 HPAIV strain (clade 2.3.4.4b, avian origin, from now on **H5N1**). These viruses were selected due to their wide and distinct host ranges, as well as the availability of *in vivo* infection rate data for some of the species included in this study^27–30,48,49^. For both pH1N1 and H5N1, we used the same experimental protocol (**Figure 2A**). We infected near-confluent 2D-monolayers with either pH1N1 or H5N1 at a multiplicity of infection between 0.01 and 0.05 and fixed replicate plates at 1-, 2-, and 3-days post-infection (dpi). We then used immunofluorescence staining of cell nuclei and influenza nucleoprotein combined with confocal imaging to quantify relative cell viability and the fraction of infected cells (see schematic of analytical approach in **Supplementary Figure 1**).

**Figure 2.**
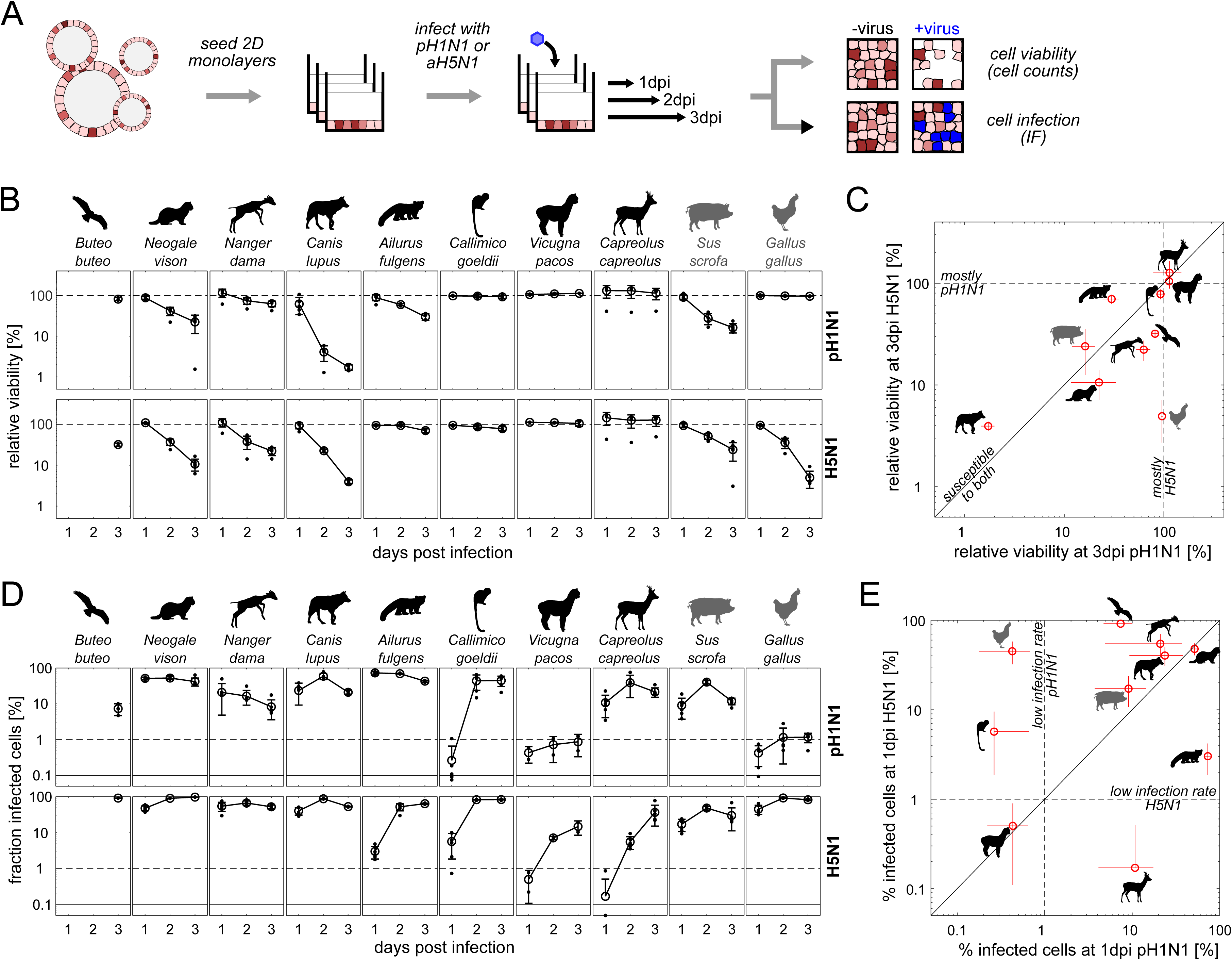
*In vitro* infection of airway organoids from wildlife and livestock species with two Influenza A viruses. **A)** Schematic of the experimental approach. **B)** Cell viability (relative to uninfected controls) during H1N1 (top) or H5N1 (bottom) infection. Large circles denote the mean across all replicates. Error bars denote standard deviation (n = 2-6). Small circles denote individual replicates. Note that some animal-timepoint combinations could not be performed due to limiting cell material during seeding. **C)** Direct comparison of relative viability between pH1N1 and H5N1 infections at 3 dpi, plotted for each animal species to illustrate differential responses across models. **D)** Fraction of infected cells during pH1N1 (top) or H5N1 (bottom) infection, as determined by IF staining of viral antigen in the cell nucleus. Large circles denote the mean across all replicates. Error bars denote standard deviation (n = 2-6). Small circles denote individual replicates. Note that some animal-timepoint combinations could not be performed due to limiting cell material during seeding. Continuous horizontal lines: detection limit at 0.1% infected cells. Dashed horizontal line: visual aid to denote threshold for low infection rate, set to <1% infected cells. **E)** Comparison of infection rates between pH1N1 and H5N1 at 1 dpi, plotted for each animal species to highlight interspecies differences in susceptibility. Note that for *B. buteo* data are shown for 3 dpi because the limiting cell material prevented seeding for 1dpi and 2dpi replicates.

First, we focused on cell viability as a primary readout of viral cytopathogenicity. Cell viability varied dramatically across animal species and between viruses (**Figure 2B-C**). For example, Goeldi’s monkey and alpaca cultures showed no apparent changes in cell viability in presence of either virus, whereas Iberian wolf culture viability dropped below 10% at 3 dpi for both viruses. Importantly, the data also recapitulated previously reported *in vivo* viral susceptibility for the two livestock species tested here: chicken culture viability was much more strongly affected by H5N1 infection^30^, whereas pig culture viability was similarly affected by both viruses (**Figure 2C**), as demonstrated before^28,29^.

Second, we focused on the fraction of infected cells (termed here infection rate), as determined by immunofluorescence staining of influenza nucleoprotein (**Figure 2D-E**). In line with the established influenza replication cycle, the viral antigen staining was in all cases largely restricted to the nucleus (**Supplementary Figure 2**), suggesting the absence of atypical infection routes among the tested species. Moreover, and similarly to the cell viability data mentioned above, the fraction of infected cells varied dramatically across animal species and viruses (**Figure 2D-E**), especially at the early stages (i.e. 1 dpi) of infection (**Figure 2E**). For example, upon exposure to pH1N1, several (but notably not all) mammal species exhibited substantial viral antigen presence already at 1 dpi, whereas other species (i.e. chicken and alpaca) showed minimal infection.

Direct comparison of infection rate and cytopathogenicity revealed a variety of distinct viral susceptibility patterns in these organoid cultures (**Figure 3**). Some species showed both low infection rate and cytopathogenicity (most prominently alpaca and chicken during pH1N1 infections), suggesting an overall low vulnerability to the respective virus. Other species showed high infection rate and cytopathogenicity (e.g. Iberian wolf infected with either virus, chicken and American mink during H5N1 infections), suggesting high vulnerability to the respective virus. A third subset of species showed low cytopathogenicity despite high rates of infection during the experiment (e.g. Goeldi’s monkey infected with either virus), suggesting an elevated tolerance to infection. Importantly, these patterns were highly virus-specific (i.e. the same animal species could show distinct infection rate/cytopathogenicity patterns depending on the virus). Thus, these results demonstrated that by infecting a panel of airway organoids from diverse animal species, we can unveil distinct modes of viral susceptibility that match available *in vivo* information.

**Figure 3.**
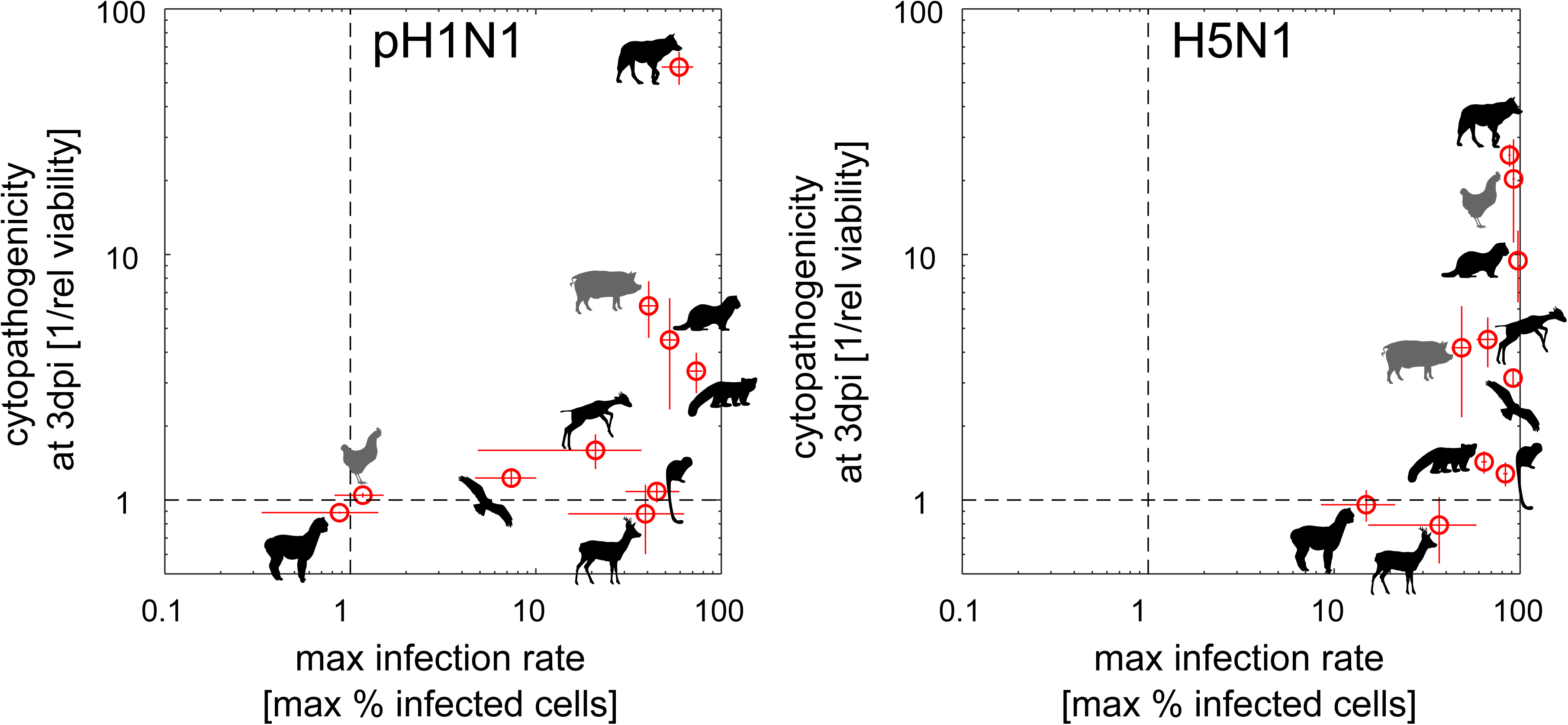
Comparative analysis of maximal infection rate and cytopathogenicity across animal organoids infected with pH1N1 (left) or H5N1 (right). Cytopathogenicity was calculated as 1/relative cell viability at 3 dpi from the data in Figure 2, while maximal infection rate was determined as the maximal % of infected cells throughout the experiment. Note that for *B. buteo*, the infection rate data are shown at 3 dpi due to technical issues because the limiting cell material prevented seeding for 1 dpi and 2 dpi replicates. Error bars denote standard deviation (n = 2-6 infected wells).

### Section 3: Host-dependent differences in influenza A receptor presence

The infection experiments described above revealed substantial interspecies variation in susceptibility to pH1N1 and H5N1 among airway organoids derived from ten different animal species. To investigate potential mechanisms underlying these differences, we next examined the expression of viral entry receptors. Influenza A viruses rely on sialic acid residues on host cell surface glycoproteins for binding and internalization. However, receptor specificity differs between strains: pH1N1 preferentially binds to α2,6-linked sialic acids, while H5N1 prefers α2,3-linked sialic acids^50^. To quantify receptor presence, we performed lectin staining on organoid-derived cells using three lectins: MAA-I and MAA-II, which detect α2,3-linked sialic acids (Siaα2-3Galβ1-4GlcNac and Siaα2-3Galβ1-3GalNac, respectively); and SNA, which detects α2,6-linked sialic acids (Neu5Acα2-6Gal). We then quantified lectin staining intensities by flow cytometry.

The data revealed substantial interspecies variation in sialic acid expression (**Figure 4A** and **Supplementary Figure 3**). MAA-I staining indicated high levels of α2,3-linked sialic acids, particularly in the avian species tested (buzzard, chicken) and in the Iberian wolf. Conversely, SNA staining was prominent in several mammalian species (e.g. American mink, red panda, Iberian wolf, Goeldi’s monkey, pig), suggesting a strong presence of α2,6-linked receptors. Control experiments using organoids derived from other chickens and pigs confirmed that these sialic acid patterns were highly conserved across individual animals from the same species (**Supplementary Figure 4**) and were consistent with known *in vivo* sialic acid expression patterns^50^.

**Figure 4.**
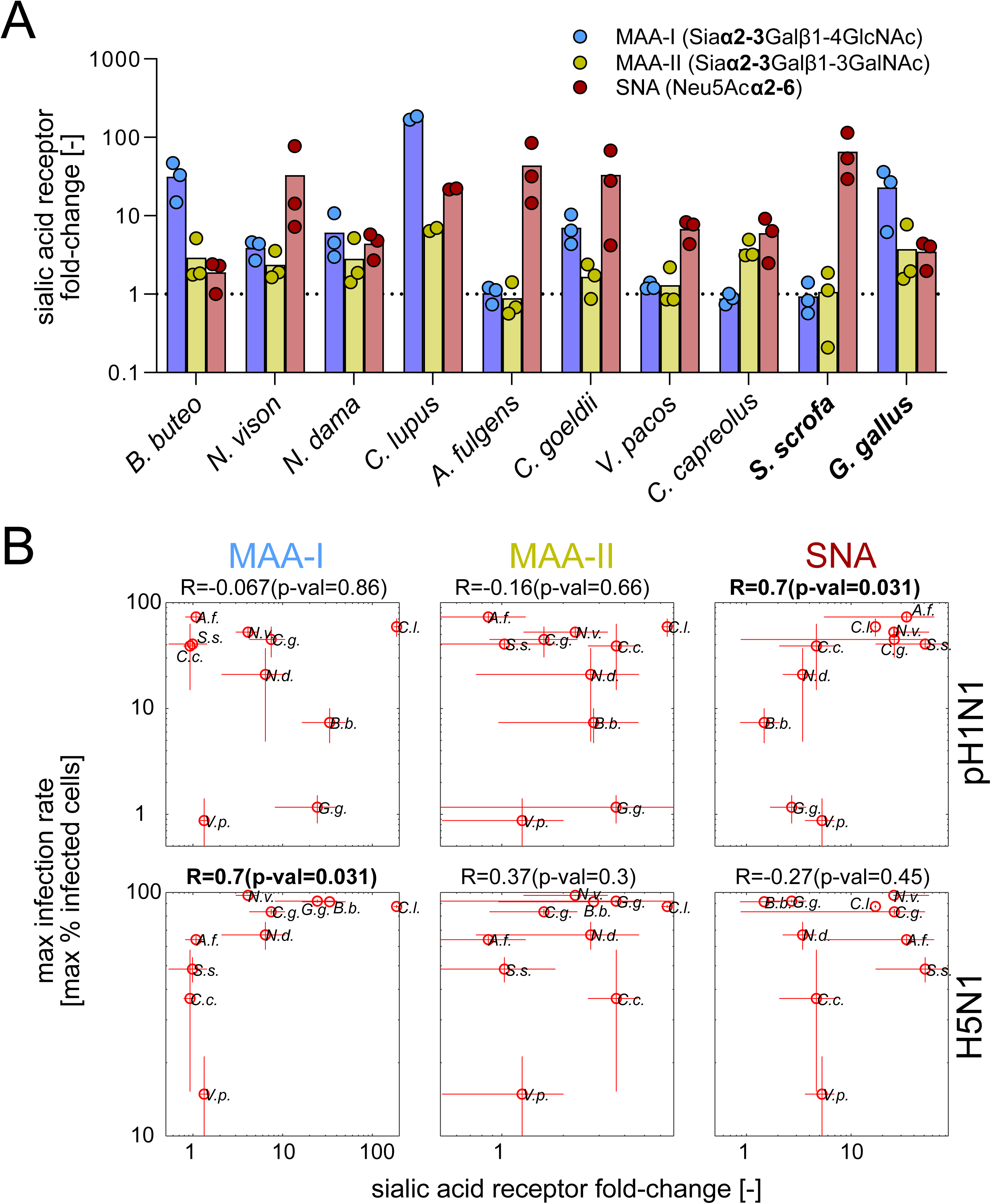
Correlation of sialic acid expression and maximal infection rate across organoids derived from different animal species. **A)** Quantification of Influenza receptor signal across animals staining for Siaα2-3Galβ1-4GlcNAc using Maackia amurensis agglutinin-I (MAA-I, blue), Siaα2-3Galβ1-3GalNAc using Maackia amurensis agglutinin-II (MAA-II, yellow), and Neu5Acα2-6 using Sambucus nigra agglutinin (SNA, red). Data shown is signal intensity as determined by flow cytometry (see methods) relative to unstained controls. Circles denote individual replicates, bars denote the mean. **B)** Same data as in A), but plotted against the maximal infection rate (calculated as the maximal % of infected cells throughout the experiment, see Figure 2) in organoid cultures infected with pH1N1 (top) or H5N1 (bottom). In each comparison, R denotes the Spearman correlation coefficient (with the respective p-value). Significant correlation coefficients (p-value < 0.05) are shown in bold. Each circle denotes an animal (designated with a two-letter code from its Latin name). Error bars denote standard deviation (n = 2-6 infected wells).

To quantitatively compare receptor sialic acid signals with previously described Influenza A infection data (Section 2), we next correlated sialic acid staining signal with the maximal infection rate (i.e. maximal % of infected cells throughout the experiment) across all animal species (**Figure 4B**). This analysis revealed some expected qualitative trends: species with high SNA signal tended to be most susceptible to pH1N1 (**Figure 4B**, top row). Conversely, species with the highest MAA-I signal, such as buzzard, chicken, Iberian wolf, tended to show the strongest infection rate upon H5N1 infection (Figure 4B, bottom row). However, even in those cases there was substantial variability across animals, suggesting that additional host-specific factors play an important role. Similarly low correlations were observed for infection rate at 1 dpi and cytopathogenicity (**Supplementary Figure 5**). Thus, these results suggest that while sialic acid presence is informative, it is insufficient on its own to explain the different viral patterns observed across the airway organoid panel.

### Section 4: Examining viral adaptation across host species

Given these large differences in influenza A virus susceptibility across this panel of animal-derived airway organoids, we next examined whether infection in these different host species could also lead to distinct patterns of viral adaptation. Our starting point towards this end was the observation that despite differences in susceptibility, none of the organoid cultures were entirely free of infected cells, particularly at later stages of infection (Figure 2D). In line with this observation, all cultures produced infectious viral particles during the experiments, pointing towards successful viral replication across species, albeit to different degrees (**Supplementary Figure 6**).

To assess whether this replication was accompanied by viral adaptation, we sequenced the viral populations at 3 dpi and compared the sequences to the original inoculum. Across both pH1N1 and H5N1 viral populations, we identified several mutations that consistently emerged in two independent replicate experiments (**Figure 5A** for pH1N1 and **Supplementary Figure 7** for H5N1). Adaptation was more pronounced in pH1N1 (**Figure 5A**), where multiple mutations were detected in the HA segment, critical for host receptor binding. Notably, a C➔A substitution at position HA-644 became the dominant sequence variant (allele frequency > 75%) in both replicates of the Goeldi’s monkey cultures.

**Figure 5.**
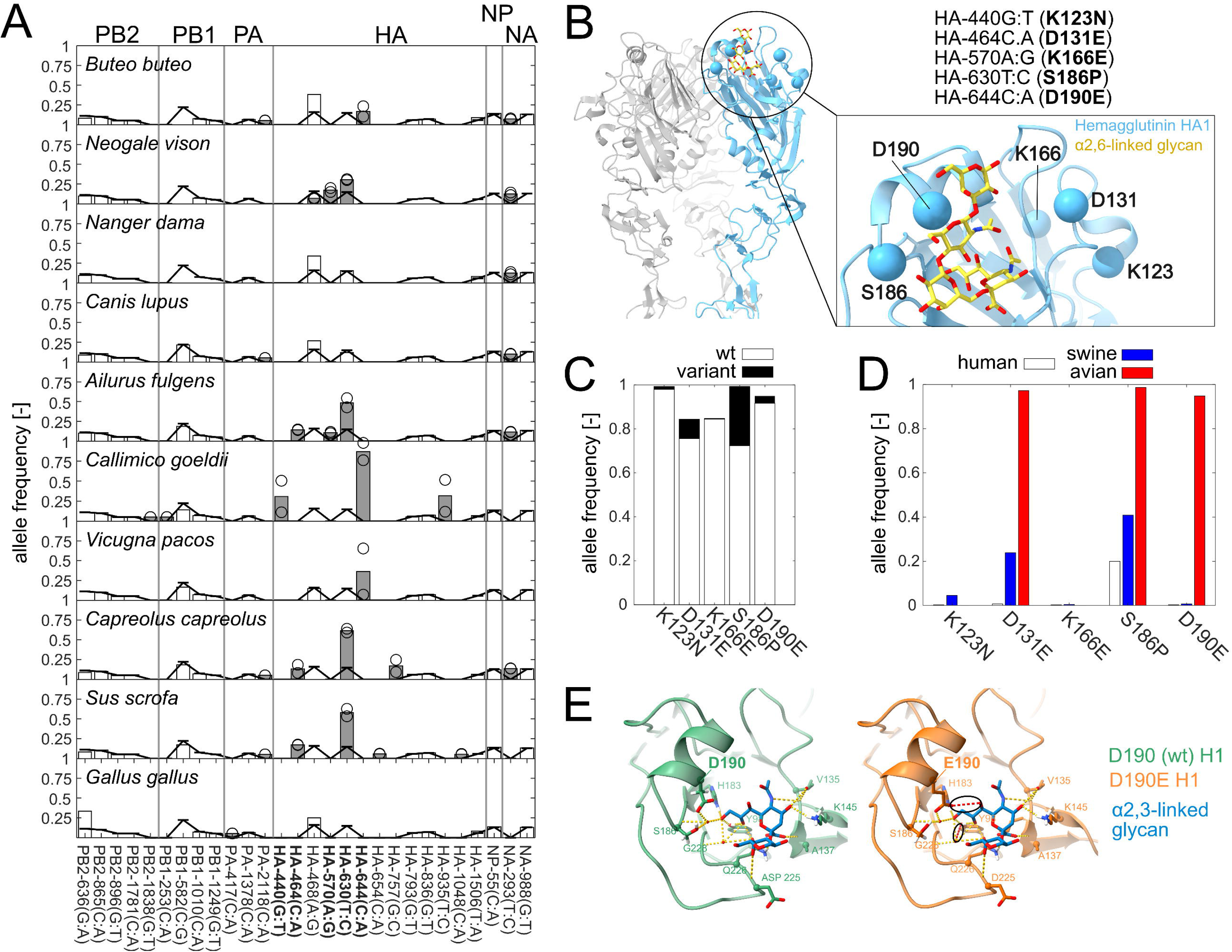
Examining viral adaptation of different organoid cultures infected with pH1N1. **A)** Analysis of mutations with changes in allele frequency in pH1N1. Only mutations identified as intra-host single-nucleotide variants (iSVNs) in two independent replicate experiments with a >2-fold difference in allele frequency compared to the inoculum are shown. Rows denote animal species, columns denote mutations (sorted based on segment and position). Continuous black line: allele frequency in inoculum. Gray bars: mutations with >2-fold difference in mean allele frequency compared to the inoculum in both replicates. Black circles: corresponding individual replicates. Mutations in bold: hotspot mutations used in subsequent panels. **B)** Protein structure of pH1N1 HA segment (PDB ID: 3UBE), visualized with ChimeraX^52^. The HA trimer is displayed in ribbon representation in blue, and the glycan ligand is shown as sticks. Mutated positions examined in the study are rendered as spheres, and labels follow the H3 numbering standard^53^. The inset table shows the mapping of genomic position (panel A) and protein position for each mutation. **C)** Prevalence of hotspot mutations in the H1N1 HA segment among organoid infection samples compared to 9,802 virus isolates from the NCBI influenza virus resource^51^. White bars: wildtype allele (wt). Black bars: mutant allele identified (see B). **D)** Same data as in panel A, stratified by host origin (human, swine, and avian only). **E)** Superimposition of experimentally resolved glycan-bound HA structure (here: alpha2,3-sialic acid) with AlphaFold2 models without (left) or with (right) the D190E substitution (see methods). Hydrogen bonds and ligand contacts were identified and visualized in ChimeraX, allowing direct comparison of how the D190E mutation influences the local interaction network for each glycan type. Black circles: visual aid showing two hydrogen bonds between HA and alpha2,3-sialic acid, one of which is water-mediated, present only with the mutated Glu190 HA.

Next, we examined the mutations found in the pH1N1-infected organoid cultures in more detail. Specifically, we focused on five mutations in the HA segment (440G➔T, 464C➔A, 570A➔G 630C➔T, 644C➔A) with low allele frequencies in the inoculum and moderate to high allele frequencies in at least one of the organoid cultures. To assess the potential functional relevance of these mutations, we mapped them to the HA protein structure. We found that all five mutations lead to amino acid substitutions in the host receptor binding domain of HA (**Figure 5B**), suggesting a potential role in receptor binding specificity. To elucidate this role, we first checked the prevalence of these mutations by examining nearly 10,000 pH1N1 hemagglutinin sequences from diverse host origins retrieved from the NCBI influenza virus resource^51^. We detected all five mutations in the database (suggesting that these mutations are not unique to the *in vitro* conditions tested here), but generally they tended to have low allele frequencies (**Figure 5C**). Nevertheless, three of these mutations were highly enriched in virus isolates from avian sources (**Figure 5D**). Consistently, our structural analyses of pH1N1 HA-sialic acid complexes suggested that the mutation D190E (i.e. 644C➔A at the genomic level) can enable the formation of two additional hydrogen bonds with α2,3-linked sialic acids. One of these bonds is formed directly between Glu190 and the sialic acid, while the other is established between Try98 and a water molecule, which in turn also interacts with the sialic acid (**Figure 5E**), suggesting a potential enhanced compatibility with α2,3-linked glycans. Taken together, these data suggest that in this organoid panel, pH1N1 undergoes rapid adaptation that can lead to highly species-dependent changes in its receptor binding specificity.

## Discussion

In this study, we aimed to develop an organoid-based approach to study the susceptibility of wildlife species to viral infections *in vitro*. To this end, we generated a panel of airway organoids from ten diverse animal species, including critically endangered ones (Table 1). We characterized the organoids and, as proof of concept, we tested their susceptibility to two influenza viruses, pH1N1 and H5N1, using three key infection metrics: cytopathogenicity, infection rate, and viral adaptation. These efforts yielded three key insights.

First, we found that a single, generalized protocol was sufficient to generate airway organoids from a diverse panel of animal species, including both mammals and birds. This is in line with previous studies that successfully applied human-optimized protocols to generate organoids in other species^32,36,39^. Importantly, our organoid panel recapitulated known *in vivo* patterns of sialic acid receptor expression —indicating that species-specific molecular features of airway tissues are retained— and were conserved within the same species in those cases we could examine (**Supplementary Figure 4**). Future studies could leverage this resource to examine whether other tissue features of interest, such as developmental programming^54^, are similarly conserved across species. One caveat to this study is that while the selection of animal species tested here is diverse, it is by no means exhaustive, and it is conceivable that more distantly related animal species, such as reptiles, amphibians, or fish, may require customized protocols for organoid generation^55,56^. Therefore, future efforts will be needed to examine the broader applicability and limitations of the approach presented here.

Second, we observed significant interspecies differences in susceptibility to influenza infection. These differences largely recapitulate known *in vivo* data for species such as pig and chicken, whose *in vitro* susceptibility was consistent with reported *in vivo* data^27–30^. However, susceptibility was only moderately correlated with sialic acid receptor abundance (**Figure 4**), highlighting the importance of further identifying host-specific factors that modulate viral infection. Notably, we identified several wildlife species with high *in vitro* susceptibility to either H5N1, pH1N1, or both, for which corresponding *in vivo* data are not available. For example, organoid cultures derived from the dama gazelle and Iberian wolf were highly susceptible to both viruses, while those from the red panda showed strong susceptibility to pH1N1. Interestingly, recent epidemiological data have shown that wild carnivore species are highly susceptible to influenza A infection^57–59^. To our knowledge, this is the first *in vitro* examination of influenza susceptibility in these species, and the information presented could support pandemic preparedness and conservation efforts by assessing the risk of future influenza outbreaks in wild or captive animal populations.

Our third insight is that none of the tested species appear to be fully resistant to influenza infection. For example, even the species with the lowest cytopathogenicity to both influenza viruses, such as alpaca and Goeldi’s monkey, produced substantial amounts of infectious viral particles within 3 dpi (**Supplementary Figure 6**). This finding is consistent with recent work demonstrating that viral entry is a weak barrier to zoonosis *in vitro*^60^, but it also provides the virus with more opportunities for adapting to a new host. In fact, in follow-up experiments, we did identify several mutations in the receptor binding domain of pH1N1 that emerge within the first 3 days of infection in a highly species-dependent manner (**Figure 5**). The most striking of these was the D190E mutation in HA segment, which became the dominant allele in Goeldi’s monkey organoid cultures in two independent infection experiments. Seminal work has shown that residue 190 is a key determinant of receptor-binding specificity in H1 hemagglutinin, and that the D190E substitution can enhance compatibility with α2,3-linked sialic acid conformations or contribute to shifts toward more avian-like receptor usage ^61^, causing a shift in host affinity towards avian species. Consistently, our structural modelling indicates that the D190E variant can form additional hydrogen bonds with α2,3-linked glycans (**Figure 5E**). Notably, our sialic acid quantification (**Figure 4**) showed that Goeldi’s monkey organoid cultures also had substantially higher α2,3-linked sialic acid signal than the other tested mammal species (with exception of the Iberian wolf). It is tempting to speculate that Goeldi’s monkey (and perhaps other, related, species with similar sialic acid expression profiles) could therefore represent a mammal host that serves as a “gateway” for pH1N1 towards avian hosts. Future studies may leverage the organoid panel presented here as a starting point to examine viral adaptation to new host species in more detail.

This study has several limitations. First, the opportunistic nature of our animal tissue sampling (i.e., we received tissue samples from animals that were euthanized for medical reasons) made it difficult to generate a comprehensive panel of animal species representing all important fauna in a particular environment of interest. Moreover, since our organoid panel included only one individual animal per species (due to restrictions in tissue sample access), we were unable to assess how variable influenza susceptibility is across individuals of the same wildlife species. Nevertheless, we note that this organoid panel can be readily expanded as more tissue samples become available. For example, since completing this study, we have successfully generated airway organoids from >20 additional species, including macropods (wallaby), sea mammals (sea lion), reptiles (iguana), big cats (Sumatran tiger), and elephants (African elephant), enabling future studies to examine viral host tropism across an even broader range of species. Similarly, as more samples from the same animal species become available, future efforts may also examine intra-species differences in viral susceptibility using the approaches developed here.

Second, to enable the rapid testing of viral susceptibility at high throughput, the infection experiments in this study used 2D monolayers derived from each animal’s 3D organoid cultures. Recent studies have shown that the cultivation system can have a dramatic impact on viral infection dynamics in some viruses. For example, experiments comparing the infection rate of human lung organoids by severe acute respiratory syndrome coronavirus 2 (SARS-CoV-2) and Middle East respiratory syndrome coronavirus (MERS-CoV) demonstrated differences between organoid-derived 2D monolayers (minimal infection) and Air-Liquid-Interface (ALI) cultures (highly susceptible)^46^. In line with these results, we similarly observed minimal infection of animal organoid-derived 2D monolayers to SARS-CoV-2, even in species known to be susceptible to the virus *in vivo* (**Supplementary Figure 8**). Therefore, it is conceivable that the approach we used in this study may underestimate susceptibility to infection also for other viruses that rely heavily on the cultivation system. Nevertheless, it is important to note that generating organoid-derived ALI cultures is complex (requiring additional differentiation steps that can take several weeks^62^) and poorly developed for most animal species. To our knowledge, beyond humans, ALI-cultures have only been successfully established for a few species (e.g., camelids^32^, pig^63^, cow^64^). Future studies will be needed to develop more efficient and robust protocols to establish ALI-cultures from diverse animal species.

In conclusion, we developed a panel of airway organoids from various wildlife species, including highly endangered ones. This novel resource enabled us, for the first time, to examine the *in vitro* susceptibility of these species to influenza infection, revealing large differences in infection rates, cytopathogenicity, and viral adaptation. This work provides an important new tool for studying the susceptibility of wildlife species to viral infections, and contributes to broader efforts in pandemic preparedness, wildlife conservation, and zoonotic disease research.

## Materials and Methods

### Reagents

Media and reagents used for airway organoid generation and subcultivation are provided in **Supplementary Table 1**. All probes used for immunodetection are provided in **Supplementary Table 2**, with the corresponding concentration at which they were used.

### Animal Tissue Collection and Sample Preservation

Airway and lung tissue samples were collected from various wildlife and livestock species. Wildlife samples were obtained from animals that were euthanized for welfare reasons unrelated with infections in the Barcelona Zoo, which included a dama gazelle (*Nanger dama mhorr*), red panda (*Ailurus fulgens)*, Iberian wolf (*Canis lupus signatus*), and Goeldi’s monkey (*Callimico goeldii*). Additional wildlife samples were collected from the Torreferrussa Wildlife Recovery Centre, namely an American mink (*Neogale vison*), and buzzard (*Buteo buteo*). Roe deer (*Capreolus capreolus*) tissue was collected by the Wildlife Ecopathology Service at Universitat Autònoma de Barcelona from a corpse involved in a car accident.

In addition to wildlife samples, tissues from livestock species were collected from control animals that were euthanized at experimental endpoints for other procedures. These included lung tissue from pig (*Sus scorfa domesticus*) and chicken (*Gallus gallus domesticus*), and tracheal samples from alpaca (*Vicugna pacos*). All tissue samples were preserved in cold AdDF++(+++) medium until further processing to ensure optimal preservation. Detailed information about each animal sample used here is provided in **Table 1**.

### Viruses used in this study

Two A influenza viruses were used in this study: pH1N1, a pandemic swine-origin A/H1N1 (A/Catalonia/63/2009) strain isolated in 2009 from a patient at Hospital Clínic in Barcelona (GenBank GQ464405-GQ464411 and GQ168897)^29^; and H5N1 HPAIV (A/Larus ridibundus/Spain/CR4062/2023) isolated from a black-headed gull (*Chroicocephalus ridibundus*) in 2023 via a passive surveillance program near the Segre river in Lleida, Spain (EPI_ISL_18983379). Both viruses were propagated at 37 °C in the allantoid fluid of 11-day-old embryonated chicken eggs from a specific-pathogen-free flock, and the infectious virus titer was determined in Madin-Darby Canine Kidney (MDCK, ATCC CCL-34) cells and measured as tissue culture infectious doses 50% (TCID50) by following the Reed and Muench method^65^.

### 3D Organoid generation and subcultivation

The organoid generation procedure was standardi<u>z</u>ed for all species and adapted from previously established protocols for human organoid production^34,41^. Briefly, lung fragments were minced and/or mucosa samples from the airways were scrapped with a scalpel at 4 °C. The disaggregated tissue was then washed and enzymatically digested following a two-step procedure. In the first step, the tissue was incubated in AdDF++(+++), supplemented with 1 mg/mL Collagenase A (C9407-25MG, Vidrafoc), for 1h at 37 °C with orbital shaking at 200 rpm. In the second step, the tissue was further digested using TrypLE (12605-028, Gibco) for 10 min at 37 °C in a water bath. After digestion, the reaction was stopped by adding chilled AdDF++(+++), and the suspension was filtered through a 70 µm cell strainer (PLC9370, Labclinics). The resulting cell suspension was pelleted by centrifugation 300 rcf for 5 min at 4 °C. The cell pellet was resuspended with 100-150 µl of ice-cold Matrigel® (45356231, Corning) and plated in 10 µl drops (4-5 droplets per well) on a 24-well plate, which were incubated as hanging droplets for 30 min at 37 °C to allow jellification. After jellification, 500 µl of antibiotic- and antimycotic-supplemented lung medium ^41^ was added to each well, and the plate was incubated at 37 °C, 5% CO2 and 95% relative humidity.

Organoids were subcultured every 7 days, with media being replaced 3-4 days after seeding. Antibiotic and antimycotic supplements were removed after the second passage, and fibroblast contamination was removed by diluting them out. For subculturing, Matrigel® domes were gently scraped and sheared with a P1000 micropipette, followed by centrifugation, and enzymatic digestion with TrypLE for 2 minutes at 37 °C. Digestion was stopped with chilled AdDF++, and the organoids were pelleted for reseeding in Matrigel® droplets. Low-passage organoids were cryopreserved in Cryostor® CS10 (100-1061, STEMCELL) for long-term storage in liquid nitrogen.

### Generation of organoid-derived 2D-monolayers

Organoid derived 2D-monolayers were seeded in PhenoPlate™ 96-well microplates (6055302, Revvity) pre-coated with 0.5% Matrigel® as follows: 3D organoids were trypsinised with TrypLE for 7 mins until reaching a single cell suspension, washed, and resuspended with Lung medium. Cells were counted with a Corning® automatic cell counter and organoids were seeded in 100 µL at a density of 20,000 cells/well for all animals except roe deer (which was seeded at 30,000 cells/well), and incubated at 37 °C, 5% CO_2_ and 95% relative humidity. After three days, 75 µL of lung medium was added to each well. Upon reaching confluence 3 days later, and 1 day before infection, media was replaced with 90 µL of fresh lung medium.

### Detection of sialic acid residues on the surface of organoid-derived cells by lectin staining

Sialic acid residues on the surface of organoid-derived cells were quantified by flow cytometry using fluorescently labelled lectins (VectorLabs): FITC-conjugated MAA-I (FL-1311-2) for Siaα2-3Galβ1-4GlcNAc, biotin-conjugated MAA-II (B-1265) coupled with streptavidin-BV515 (STAR210SBV515, BioRad) for Siaα2-3Galβ1-3GalNAc, and Cy5-conjugated SNA (CL-1305-1) for Neu5Acα2-6. For each animal species, a single-cell suspension was prepared from 3D organoid cultures by incubation with TrypLE at 37 °C for at least 10 min. Digestion was stopped with chilled FACS buffer (PBS + 1% FBS) and cells were washed with PBS. Afterwards, cells were counted with Corning® automated cell counter, and 250.000 cells/well were plated in a U-bottom 96 well-plate (650101, Greiner Bio-One). Viability staining was performed with the LIVE/DEAD Fixable Violet Dead Cell Stain (L34955, ThermoFisher Scientific) at 1/8000 dilution in PBS for 15 min at room temperature in the dark. Cells were washed twice with PBS and incubated with all lectins at a concentration of 1 µg/mL in FACS Buffer for 1 hour at room temperature in the dark. Then, cells were washed twice again and incubated with the streptavidin-SBV515 secondary probe for 20 min on ice in the dark. Finally, cells were washed twice again with PBS and fixed with 1% paraformaldehyde (PFA) and run in a MACSQuant® Analyzer (Miltenyi Biotec). The analysis was performed using FlowJo v10.10.0 (Tree Star) using a gating strategy as shown in **Supplementary Figure 3**.

### Viral infection of organoid-derived 2D-monolayers

Organoid infection with the aforementioned Influenza A virus strains was performed in BSL-3 facilities under the approval of IRTA-CReSA’s Biosafety Committee (CBS 159-2024) as follows: organoid-derived 2D monolayers that were seeded in 96-well plate format 1 week prior (see above), and whose media was replaced with 90 µL/well Lung medium one day before infection, were infected with either pH1N1 or aH5N1. Infections were performed by addition of 10 µL of the viral preparation (5714 TCID_50_, corresponding to a multiplicity of infection of 0.01-0.05) to each culture. Cells were incubated for up to 3 days at 37 °C, 5% CO_2_ and 95% relative humidity. On days 1, 2, and 3 post-infection (dpi), supernatants were recovered for viral quantification (RT-qPCR and titration), and cells were fixed as described in the section “*Immunofluorescence staining of organoid-derived 2D monolayers*”.

### Immunofluorescence staining of organoid-derived 2D monolayers

Cultures in 96-well plate format were fixed for immunofluorescence staining as follows: the medium of each well was aspirated, and the monolayers were fixed with 4% PFA in PBS (50µL/well) for 10 minutes at RT (30 min in the case of samples infected with viruses). PFA was aspirated, and the wells were washed 3 times with PBS (5 minutes at RT each wash). The plates were sealed with parafilm and stored at 4 °C until use.

For the immunostaining of the fixed samples, the PBS was aspirated, and the samples were permeabilized with 1% Triton™ X-100 in PBS for 1h at RT (50µL/well). The samples were then washed 3 times with 0.1% Tween® 20 in PBS (5 minutes at RT each wash, 100µL/well) and incubated with Blocking buffer (PBS supplemented with 10% FBS and 0.5% Triton™ X-100, filtered 0.2µm) for 1h at RT (30 µL/well). The blocking buffer was aspirated, and the samples were incubated for 24h at 4 °C with the primary antibodies (30 µL/well) diluted in Dako REAL antibody diluent (52022, Agilent) at the concentrations specified in **Supplementary Table 2**. The samples were then washed 5 times with wash buffer (5 minutes at RT each wash, 100µL/well) and incubated for 24h at 4 °C with the secondary antibodies (30 µL/well) diluted 1/400 in Dako REAL antibody diluent at the concentrations specified in the key resources table. Hoechst 33342 (4µg/mL) or Phalloidin Atto 488 (1/500) were added together with the secondary antibodies. The samples were washed 3 times with 0.1% Tween® 20 in PBS and 2 twice with PBS alone (5 minutes at RT each wash, 100µL/well). Finally, 150µL of 0.02% Sodium azyde in PBS was added per well and the plates were stored at 4 °C sealed with parafilm until imaging. All the incubations steps starting from permeabilization were performed under mild agitation (50 rpm) in a humidified chamber.

### Image acquisition

3D organoids inside Matrigel domes (Figure 1B) and videos of cilia movement in 3D and 2D (**Supplementary videos 1** and **2**) were imaged with an Olympus CKX53 microscope with a 4x objective model UPLFLN4XIPC-2 (Figure 1B) or a 20x objective model LCACHN20XIPC (**Supplementary video 1**) or a 10x objective model CACHN10XIPC (**Supplementary video 2**, left) or a 40x objective model LCACHN20XIPC (**Supplementary video 2**, right zoom).

Immunostained organoid-derived monolayers were imaged with a Leica Stellaris 8 confocal microscope with a HC PL APO 10x/0,40 CS2 dry objective (#11506424) (Supplementary Figures 1 and 2) or a HC PL APO 40x/1.30 OIL CS2 objective (#11506428) (Figure 1C). For each image, a multidimensional stack was acquired at 2048×2048 pixel resolution (pixel size = 0.57 µm for 10x and 0.14 µm for 40x). Stacks comprised multiple z-planes covering the full height of the monolayer with a total z-range of 40µm, with 5 planes spaced 10µm and 80 planes spaced 0.5µm for 10x and 40x objectives, respectively, including multiple fluorescence channels for each dye. To image the full area of each well of the 96 well plates (**Supplementary Figure 1**), overlapping tiles were acquired at 10% overlapping area with the 10x objective. The overlapping tiles were then merged with the Leica Application Suite X software (v4.8.0.8989) into a single image stack. For presentation purposes, representative images were further processed in ImageJ^66^. This processing included maximum intensity projection, contrast adjustment, pseudo-coloring, cropping or scalebar addition.

### Quantification of the relative viability of influenza-infected organoid cultures

To quantify the relative viability of influenza-infected organoid-derived monolayers, we used the merged multidimensional stacks covering the full wells of the 96-well plates and a custom-made image analysis pipeline coded in python (see **Supplementary Figure 1A**). For each well, a maximum intensity projection (MIP) of the nuclear staining was performed. The MIP was then percentile-normalized. This normalization was based on the distribution of signal intensity within the original image and resulted in a normalized image with pixel values in the range 0-1, where pixels with values below the percentile 0.001 in the original image were set to 0 and pixels with values above the percentile 99.999 in the original image were set to 1.

The nuclei in the normalized MIP were then segmented using Cellpose3^67^. First, the image restoration model “deblur_nuclei” was applied to the normalized MIP. Then, the segmentation model “nuclei” was applied to the restored image with the following settings: diameter= 12, normalize = False, flow_threshold= 0, cellprob_threshold = -3.

To eliminate spurious segmentation artifacts (small segmented objects that are not nuclei), the nuclear masks were filtered out based on their size. For each animal species, the distribution of nuclear mask sizes of all the control wells were combined into a single distribution. Then, the percentile 1 of this control distribution was calculated and used as a threshold to filter out the nuclear masks of all the wells of the same animal species (both control and infected wells). For each well, the nuclear masks with a size lower than the calculated threshold were filtered out. Once filtered, the number of nuclear masks in each well was determined, and the relative viability was calculated for each infected well of each animal species following the formula:

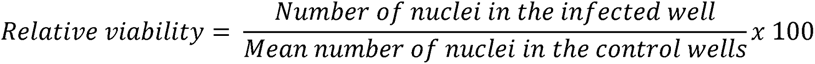

### Quantification of the percentage of infected cells in influenza-infected organoids

The percentage of infected cells was determined with a custom-made image analysis pipeline coded in python, by using the segmented and filtered nuclei segmentation masks and the signal of the nucleoprotein staining (see **Supplementary Figure 1B**). For each well, a maximum intensity projection (MIP) of the influenza nucleoprotein staining was performed. Then, the nuclear masks of the same well (detected and filtered as explained in the section “Quantification of the relative viability of influenza-infected organoids”) were used to quantify the integrated signal intensity of the nucleoprotein staining inside each nucleus. For each animal species, the distribution of integrated nucleoprotein intensity inside the nuclei of all the control wells were combined into a single distribution. Then, the percentile 99.99 of this control distribution was calculated and used as a threshold to detect the influenza-positive nuclei of all the wells of the same animal species (both control and infected wells). Those nuclei whose integrated density for influenza nucleoprotein was higher than the threshold were classified as positive (infected).

Closer examination revealed instances of non-infected cells with a moderate integrated density of the influenza nucleoprotein (above the threshold) that was caused by neighboring infected cells with a very bright nucleoprotein signal. In these false-positive cells, the nucleoprotein signal was not homogeneous within the nucleus, with higher intensity values near the true-positive neighboring cells. To filter out such false-positive cells, the nuclear centroid and the weighted nuclear centroid (centroid of the nuclear mask weighted by the distribution of the nucleoprotein intensity within the mask) were calculated with the function “regionprops_table” of the scikit-image python package^68^. The Euclidean distance between the centroid and the weighted centroid of each nucleus was calculated. Then, the nuclei showing a Euclidean distance higher than 3.5 were filtered out and classified as negative (non-infected). Finally, the percentage of infected cells in each well was calculated following the formula:

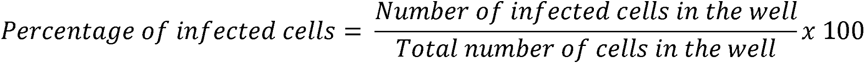

### Quantification of replicative viral particles

Quantification of replicative viral particles in the supernatants of infected organoid-derived 2D-monolayers was performed in MDCK cells. Briefly, MDCK cells were seeded at 30.000 cells/well in a 96-well plate the day before titration and incubated at 37 °C, 5% CO2, and 95% relative humidity. After 24 h, the medium was aspirated, the cells were washed with PBS (for pH1N1) and 50 µL of serially diluted supernatants (1/10) from 10^-1^ to 10^-10^ in titration medium (See supplementary Table 1) –with or without TPCK-trypsin (T8802, Merck), for pH1N1 or H5N1, respectively– were added to the respective wells (8 replicas per dilution, 16 cell controls). One hour after, 100 µL of titration medium - with or without TPCK-trypsin as indicated before- was added per well. Three days after, cytopathic effect was assessed, and viral titers were calculated in TCID50/mL ^65^.

### Influenza A RT-PCR

Viral RNA extraction from the supernatants of organoid monolayers was performed using a MagAttract 96 Cador Pathogen Kit (509947457, Indical Biosciences) in a QIAGEN BioSprint 96 Purification system (QIAGEN), following producer’s specifications. First, supernatants were inactivated with VXL buffer, followed by automated RNA purification as specified by manufacturers. RT-qPCR was performed with 3 µL of extracted RNA using an AgPath-ID™ One-Step RT-PCR Reagent (4387391, Invitrogen) targeting gene M of influenza viruses (M+ 64: FAM-5’- TCAGGCCCCCTCAAAGCCGA-3’-TAMRA). The amplification protocol consisted of a pre-amplification cycle of 10 min at 48 °C and 10 min at 95 °C, followed by 40 cycles of 2 seconds (s) at 97 °C and 30 s at 61 °C in an Applied Biosystems™ 7500 Fast Real Time PCR thermocycler.

### Influenza A whole-genome sequencing and analysis

Whole-genome sequencing (WGS) of supernatants collected from organoid cultures infected with Influenza A (either H1N1 or H5N1) was performed as follows. Briefly, viral RNA was extracted from the organoid culture supernatants using the QIAamp Viral RNA Mini Kit (52904, Qiagen), followed by RNA quantification and purity assessment using a 2100 Bioanalyzer (Agilent). After confirming RNA quality, sequencing libraries were prepared using the Illumina Microbial Amplicon Prep Influenza A/B protocol (20106305, Illumina, San Diego, CA) according to the manufacturer’s instructions. Sequencing was carried out on a NextSeq 1000 platform (Illumina) with a P1 flow cell, employing 2×150 bp paired-end reads. Paired-end Illumina reads were preprocessed with fastp^69^ using fixed parameters to ensure uniform quality across samples: the first 25 bases of both R1 and R2 were trimmed (--trim_front1 25 --trim_front2 25), bases were required to meet a minimum Phred quality of 20 (--qualified_quality_phred 20), and reads shorter than 50 nt after trimming were discarded (--length_required 50).

Influenza genome analysis was performed with the nf-flu (3.1.10) workflow in Illumina mode (--platform illumina), which encompasses read mapping, small-variant calling, depth-aware masking, and per-segment consensus generation. Because the default nf-flu behavior is to select the best-match reference independently for each sample and segment, we implemented a two-stage strategy to enforce fixed references and thereby improve comparability across samples. First, we ran nf-flu only on the two baseline controls (H5N1 and pH1N1) to obtain high-quality, depth-masked consensus sequences for all eight segments of each virus. These baseline consensuses were then used as the sole references in a second set of runs that processed all remaining samples. In these fixed-reference runs we explicitly overrode reference selection by providing the baseline consensus FASTA to the pipeline (--ref_db <baseline_consensus.fasta>). To satisfy nf-flu’s inputs while preventing the retrieval or selection of alternative references, we also supplied a minimal FASTA (the same baseline consensus, optionally compressed) and a small metadata file via --ncbi_influenza_fasta and --ncbi_influenza_metadata. Samples were processed in two cohorts (H5N1 and H1N1), each against its corresponding baseline reference set. This double-run approach ensured that read mapping, variant calling, and consensus derivation for a given virus were performed against an identical per-segment reference across species and replicates, enabling direct cross-sample comparisons.

Variant calls produced by the nf-flu workflow were post-processed from VCF to retain only high-confidence sites before classifying within-host alleles. We restricted analyses to records marked PASS, which required a defined alternate-allele frequency (AF), and excluded positions with a masked reference base (REF = N). To avoid complexities of multi-allelic loci, we kept sites with a single alternate allele (NUMALT = 1). Depth and support thresholds were imposed to ensure robust evidence for the alternate allele: total depth (DP) had to be ≥ 500 reads and alternate observations (AO) ≥ 20 reads. We further required strong mapping quality for ALT-supporting reads (mean mapping quality, MQM ≥ 40), strand balance (ALT evidence present on both strands; SAF > 0 and SAR > 0), and positional balance within reads (ALT supported on both left and right sides of the variant; RPL > 1 and RPR > 1). Only variants with high model support were retained, using an odds threshold of ODDS ≥ 1000.

After this general filtering, intra-host single-nucleotide variants (iSNVs) were defined as alleles with AF >= 0.02, site depth (DP ≥ 1000) and AO ≥ 20. Moreover, only iSNVs that passed these filters in the samples from both independent experiments were considered. The sole exception was the H5N1 pig infection experiment, for which only one sample passed QC; in that case a variant was retained if detected in the single available sample.

### Analysis of mutation prevalence in influenza sequences

To determine the prevalence of the identified HA mutations in pH1N1 strains, we retrieved hemagglutinin sequences from the NCBI Influenza Virus Resource^51^. Protein sequences were filtered by influenza virus type (A), protein (HA), and subtype (H1N1), with no restrictions on host or geographic origin, and a minimum sequence length of 400 amino acids. Identical sequences were removed to eliminate redundancy. As of 17 November 2025, this yielded 9,802 unique sequences from diverse host origins. A multiple sequence alignment was generated using MAFFT v7.525^70^ with the “Auto” setting, and the resulting alignment was used to calculate the allele frequency of each mutation of interest, both overall and stratified by host species.

### Structural analysis of pH1N1 HA segment (D190 and D190E variants)

Experimentally resolved glycan-bound structures of HA were combined with AlphaFold2 models^71^ as follows: Crystal structures of the 2009 pandemic H1 hemagglutinin bound to α2,6-linked (LSTc; PDB 3UBE) and α2,3-linked (LSTa; PDB 3UBJ) glycans were used as structural templates. AlphaFold2 models of the HA consensus sequence, with and without the D190E substitution, were generated and superimposed onto the glycan-bound structures to position the modeled proteins within an experimentally validated receptor-binding context. This approach was necessary because AlphaFold models do not include bound water molecules, which are critical for accurately depicting HA–glycan interactions. Hydrogen bonds and ligand contacts were identified and visualized in ChimeraX^52^, allowing direct comparison of how the D190E mutation influences the local interaction network for each glycan type.

### SARS-CoV-2 infection and viral replication assays

Vero E6 cells (ATCC CRL-1586) were cultured in Dulbecco’s modified Eagle medium, (DMEM; Invitrogen) supplemented with 5% fetal bovine serum (FBS; Invitrogen), 100 U/mL penicillin and 100 µg/mL streptomycin (all ThermoFisher Scientific). Nanoluciferase SARS-CoV-2 clone (icSARS-CoV-2-nLuc; NR-54003) was obtained from BEI resources (NIAID; NIH; contribution from Dr. Ralph S. Baric). Viral stocks were titrated in Vero E6 cells, and TCID_50_/mL were calculated using the Reed & Muench method.

The Biosafety committee of the Research Institute Germans Trias i Pujol approved this study with reference approval CSB-25-007 performed at the BSL3 laboratory of the Centre for Comparative Medicine and Bioimage (CMCiB). Vero E6 and/or organoid derived 2D monolayers (1- 4×10^4^ cells per well) were seeded in white 96-well plates. After 24h, cells were inoculated with nanoluciferase SARS-CoV-2 (MOI 0.1 to 10.0) and treated or not with Remdesivir [25µM]. Non-inoculated cells and inoculated wells without cells were used as negative and background controls of infection, respectively. Viral-associated cytopathic effect was daily revised by light microscopy. Nanoluciferase expression was assessed 72h after SARS-CoV-2 inoculation using a Nano-Glo Luciferase Assay (Promega; #N1110) and measured with a Fluoroskan FL Plate Reader (Thermo Fisher).

## Supporting information

data set 1 - sequence variants

High-resolution version of Figure 5

## Data availability

Raw influenza sequencing data were deposited at the European Nucleotide Archive (https://www.ebi.ac.uk/ena/browser/home, accession number PRJEB104894). The corresponding processed variant data are provided as **data set 1**.

## Supporting information

Supplementary information is provided as:

- Single SI document containing Supplementary Table 1, Supplementary Table 2, the captions of Supplementary videos 1-2, and Supplementary Figures 1-8
- Supplementary video 1
- Supplementary video 2

## Acknowledgements

We thank Natàlia Majó Masferrer and Joaquim Segalés for fruitful discussions on the *in vivo* host tropism of Influenza A in wildlife species. We thank Laura Bonillo, Óscar Pérez and Alejandro Moreno for their help during the processing of some samples. We thank Xavier Abad, Mercedes Mora, Marta Valle, Laura Rigol, Sierra Espinar, Georgina Cuadriello for their support at the BSL-2 and BSL-3 facilities. We thank Mónica Pérez, Rosa López and Gema García, for their support at the Histopathology facility. We thank Àlex Cobos, Enric Vidal, Gemma Guevara and Diego Pérez for their support in some necropsies.

We dedicate this work to the memory of Dr. Hugo Fernández-Bellon, a veterinarian whose passion for research and commitment to animal welfare greatly enriched this study. His contributions and spirit will be remembered with deep gratitude.

## Author contributions

Conceived and designed the study: FTF, GC, KK, JVA. Performed experiments and analyses: FTF, GC, FCM, MC, MP, STA, EFT, RB, VA, NC, JE, HFB, LFM, RM, NN, MPS, JR, NR, OT, RV, MRD. Supervised analyses: AV, RP, NIU, MRD, KK, JVA. Wrote manuscript with contributions from all authors: FTF, GC, KK, JVA.

