## Supplementary Information for "Airway organoids reveal patterns of Influenza A tropism and adaptation in wildlife species"

### Supplementary tables

#### Supplementary table 1. Cell culture media and buffers used in this study

| **Buffers and media** | | | | |
| --- | --- | --- | --- | --- |
| Buffer & Media | Compound | Conc. | Reference | Provider |
| AdDF++ | DMEM/F-12 | 1X | 12634-010 | Gibco |
|  | Glutamax | 1% | 35050-061 | Gibco |
|  | Hepes 1M | 1% | 15630-080 | Gibco |
| AdDF++(+++) | DMEM/F-12 | 1X | 12634-010 | Gibco |
|  | Glutamax | 1% | 35050-061 | Gibco |
|  | Hepes 1M | 1% | 15630-080 | Gibco |
|  | Penicillin Streptomycin | 100 U/mL 100 µg/mL | 15140-122 | Gibco |
|  | Primocin | 50 µg/mL | Ant-pm-1 | InvivoGen |
|  | Amphotericin B | 0.5 µg/mL | 15290-026 | Gibco |
| Lung medium | N-Acetylcysteine | 1.25 mM | A9165-5g | Sigma |
|  | Nicotinamide | 5 mM | N0636 | Sigma |
|  | B27 supplement | 1X | 17504-44 | Gibco |
|  | SB202190 | 500 nM | S7067 | Sigma |
|  | A83-01 | 500 nM | 2939 | STEMCELL |
|  | Y-27632 | 5 µM | 72304 | STEMCELL |
|  | R-spondin | 500 ng/mL | 78213 | STEMCELL |
|  | Noggin | 100 ng/mL | 78060 | STEMCELL |
|  | FGF-7 | 25 ng/mL | 78046 | STEMCELL |
|  | FGF-10 | 100 ng/mL | 78037 | STEMCELL |
| Titration medium | DMEM | 1X | 10-013-CV | Corning |
|  | Glutamax | 1% | 35050-061 | Gibco |
|  | Penicillin Streptomycin | 100 U/mL 100 µg/mL | 15140-122 | Gibco |
|  | Bovine serum Albumin | 0.3% | A7906 | Sigma |
| FACS Buffer | PBS | 1X | 21-040-CV | Corning |
|  | Fetal bovine serum | 1% | 35-079-CV | Corning |
| Permeabilising buffer | PBS | 1X | 21-040-CV | Corning |
|  | Triton X-100 | 1% | T8787 | Sigma |
| Wash buffer | PBS | 1X | 21-040-CV | Corning |
|  | Tween® 20 | 0.1% | P1370 | Sigma |
| Blocking buffer | PBS | 1X | 21-040-CV | Corning |
|  | Triton X-100 | 0.5% | T8787 | Sigma |
|  | Fetal bovine serum | 10% | 35-079-CV | Corning |
| Storage buffer | PBS | 1X | 21-040-CV | Corning |
|  | Sodium azyde | 0.02% | S-8032 | Sigma |
| Fixation buffer | PBS | 1X | 21-040-CV | Corning |
|  | Paraformaldehyde | 4% | J199943.K2 | Invitrogen |
| **Other cell culture reagents** | | | | |
| Reagent | Reference | | Provider | |
| Collagenase A | C9407-25MG | | Vidrafoc | |
| TrypLE | 12605-028 | | Gibco | |
| Matrigel® | 45356231 | | Corning | |
| Cryostor®CS10 | 100-1061 | | STEMCELL | |
| TPCK-Trypsin | T8802 | | Merck | |

#### Supplementary table 2. Probes for immunodetection used in this study

| Probe | Concentration used | Reference number | Provider |
| --- | --- | --- | --- |
| FITC-conjugated MAA-I | 1 µg/mL | FL-1311-2 | VectorLabs |
| Biotin-conjugated MAA-II | 1 µg/mL | B-1265 | VectorLabs |
| Cy5-conjugated SNA | 1 µg/mL | CL-1305-1 | VectorLabs |
| Streptavidin-BV515 | 1 µg/mL | STAR210SBV515 | BioRad |
| Hoechst 33342 | 1/2500 | H3570 | Invitrogen |
| Phalloidin Atto 488 | 1/500 | 49409 | Sigma |
| Cytokeratin 5 (RCK103) | 1/200 | SC-32721 | SantaCruz |
| Purified mouse anti-Ki-67 | 1/500 | 550609 | BD |
| Purified mo anti-E-cadherin | 1/1000 | 610181 | BD |
| ZO-1 mAb (ZO1-1A12) | 1/200 | 339100 | Invitrogen |
| Goat anti-mo IgG AF555 | 1/400 | A21424 | Invitrogen |

### Supplementary videos

Supplementary video 1. Cilia movement in 3D organoids

Video of a porcine 3D lung organoid. Note the movement of the cilia of some cells at their apical surface, facing the internal organoid lumen. Cell debris inside the lumen rotate due to the flow generated by cilia movement. Scale bar = 100µm.

#### Supplementary video 2. Cilia movement in 2D organoid monolayers

Left: video at low magnification of a porcine 2D lung organoid monolayer. Scale bar = 100µm. Right: Video at high magnification of the region of the well indicated with a red square in the left video. Scale bar = 50µm. Note the movement of the cilia of some cells at their apical surface, facing the medium. Cell debris vibrates due to the cilia movement of the cells underneath.

### Supplementary figures

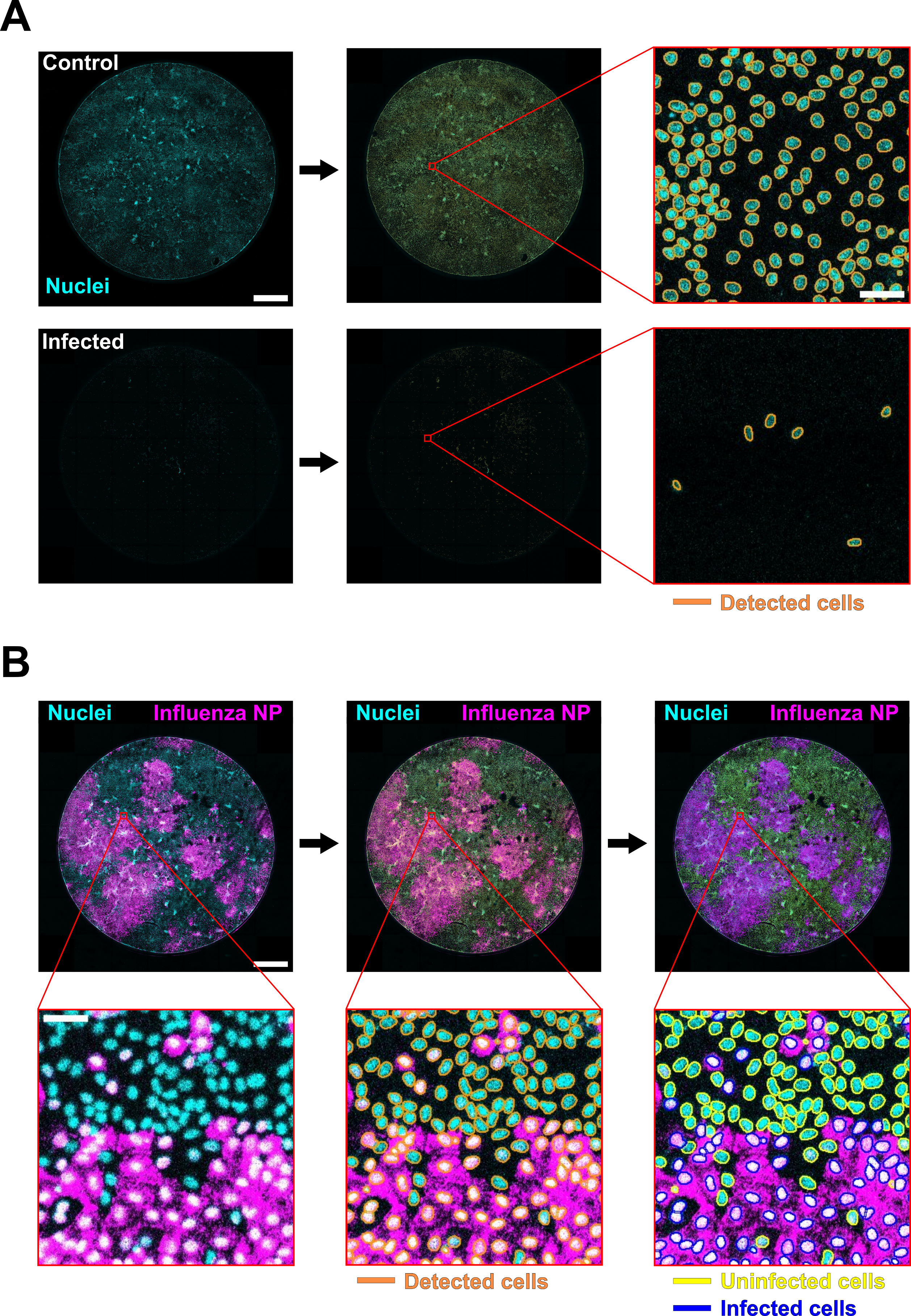

**Supplementary Figure 1. Quantification of relative cell viability and percentage of infected cells from immunofluorescence images. A)** Quantification of the relative viability by counting cell nuclei in controls and infected organoid cultures. Example shown: organoid-derived 2D-monolayers from chicken, infected with aH5N1 at 3dpi. Top: uninfected culture serving as control. Bottom: infected culture. Left: images of the nuclei staining (cyan) of a full well of a 96 well-plate. Middle: detection of cell nuclei in the left images. Right: zoomed field of views from the regions indicated with a red square in the middle images. Detected cell nuclei are outlined in orange. Scale bar of the whole-well images (left and middle) = 1mm. Scale bar of the zoomed fields of view (right) = 30µm. **B)** Quantification of the percentage of infected cells by first detecting cell nuclei (same as in A), and second by identifying nuclei with IF signal from Influenza nucleoprotein staining (see methods). Example shown: organoid-derived 2D-monolayers from chicken infected with aH5N1 at 1dpi. Top: Images of the whole-well of the 96 well-plate. Scale bar = 1mm. Bottom: zoomed field of views of the regions indicated with a red square in the top images. Scale bar = 30 µm. Left: immunostaining of nuclei (cyan) and Influenza nucleoprotein (NP). Middle: detection of cell nuclei (detected nuclei are outlined in orange). Right: classification of detected nuclei in infected or uninfected based on the NP signal (infected cells’ nuclei are outlined in blue and uninfected ones in yellow).

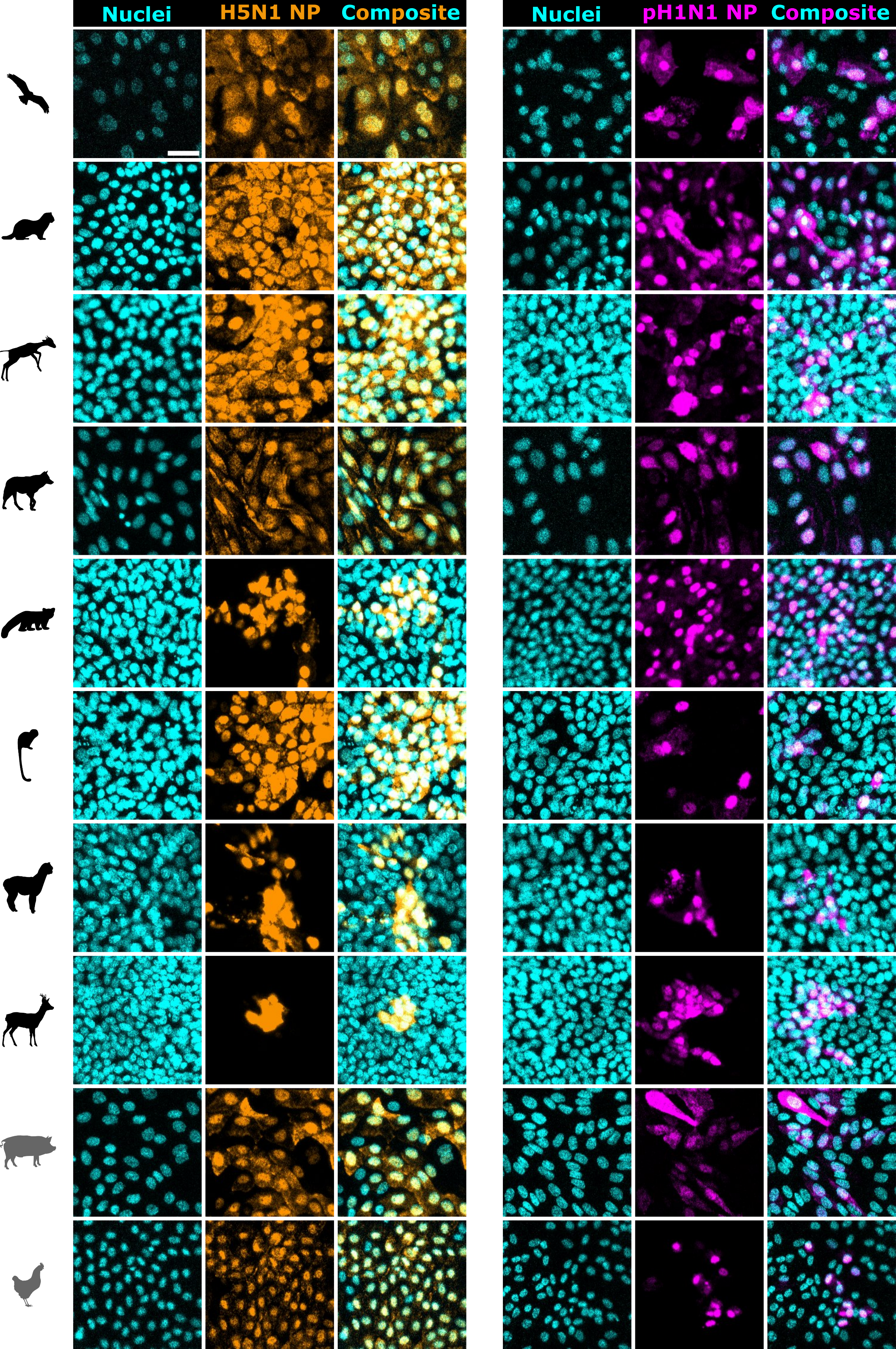

**Supplementary Figure 2. Influenza nucleoprotein distribution in infected 2D organoids.** Representative immunofluorescence images of 2D organoids infected with H5N1 or pH1N1. The cell nuclei are shown in cyan, the aH5N1 influenza nucleoprotein (NP) is shown in orange and the pH1N1 influenza nucleoprotein (NP) is shown in magenta. Each row of images corresponds to a specific animal species, indicated with a symbol at the left of each row. Scalebar = 30 µm.

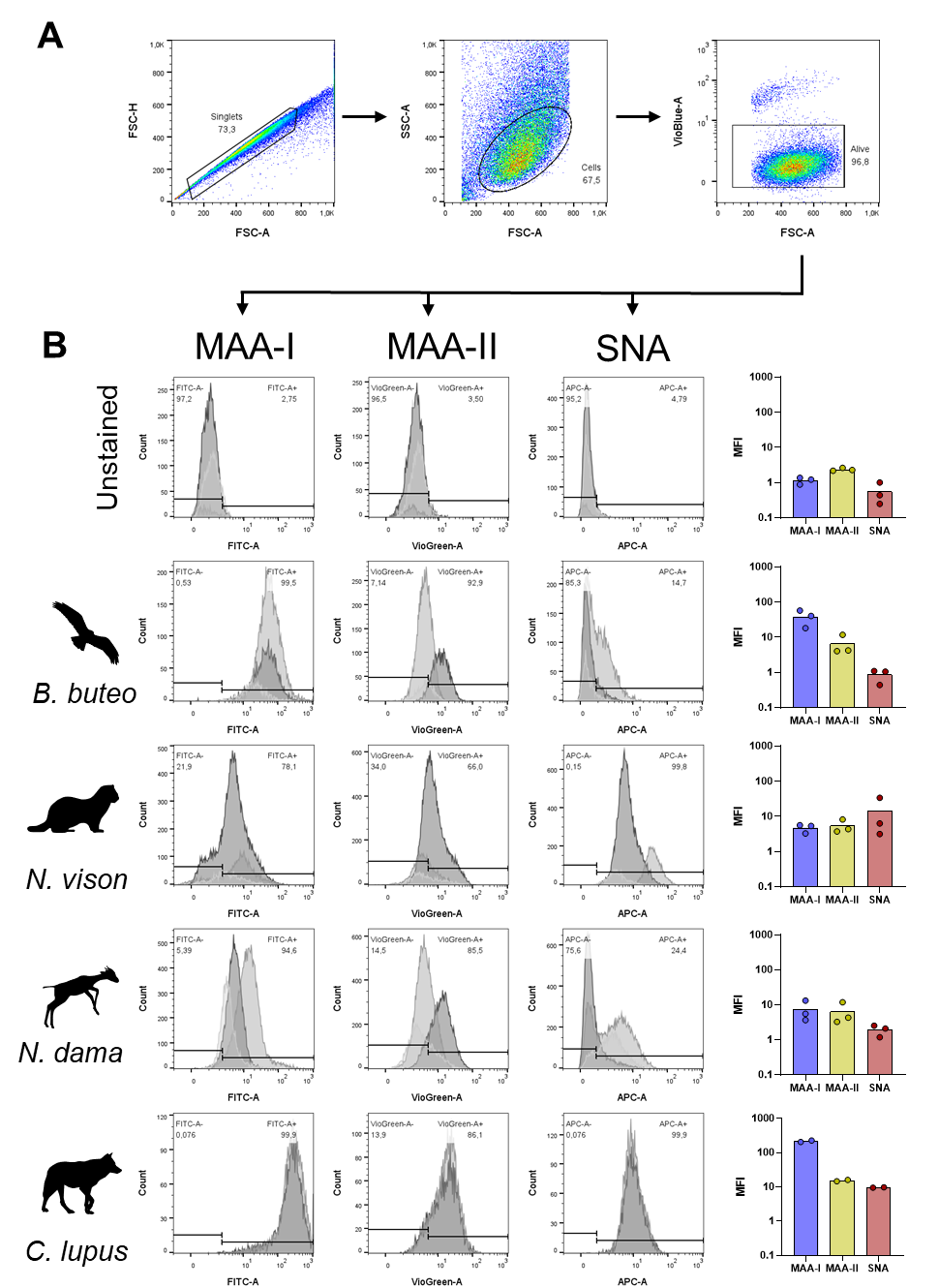

**(Supplementary Figure 3, continues on next page)**

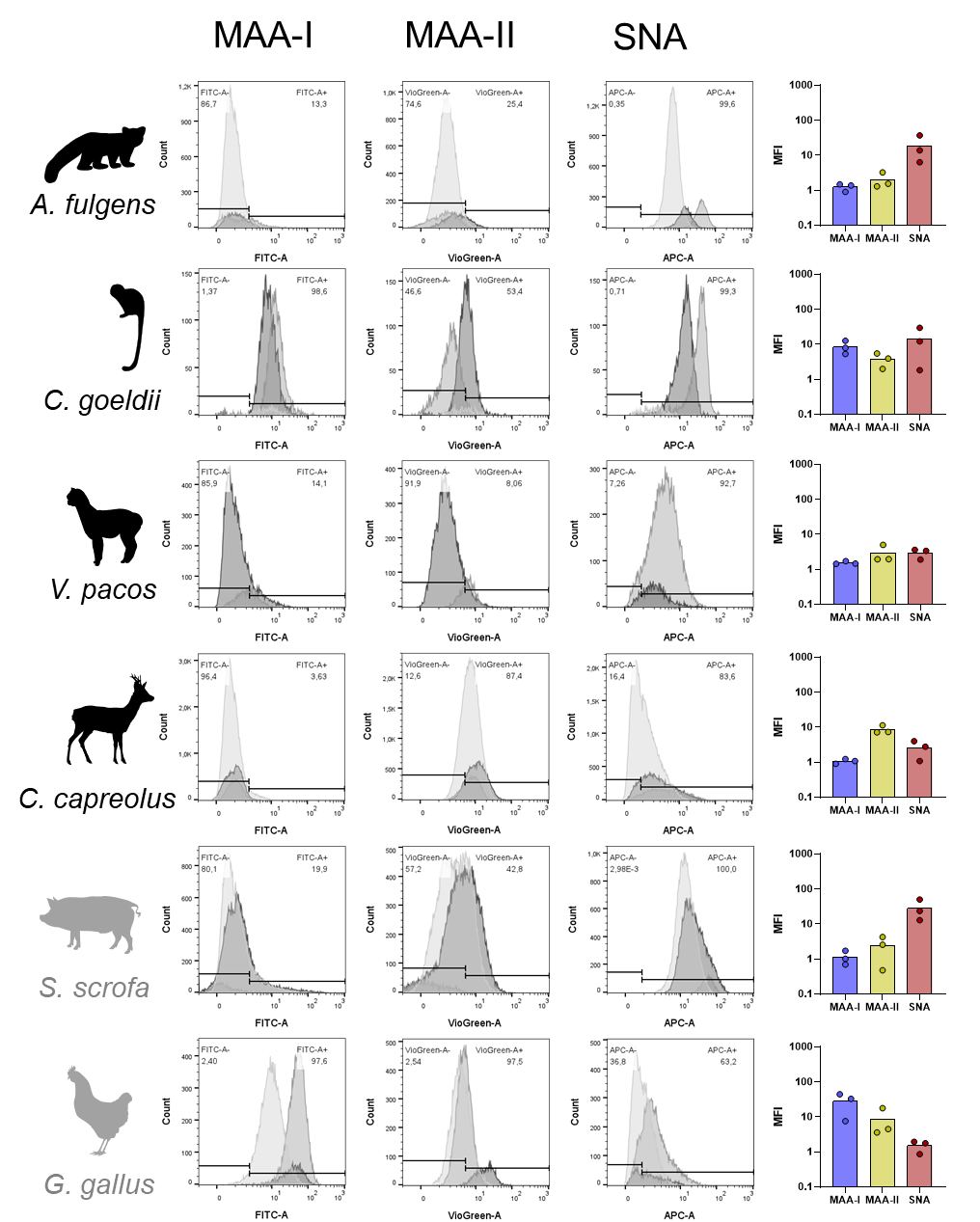

**Supplementary Figure 3. FACS analysis of wild animal-derived organoids stained with sialic acid-specific lectins.** A) Dot plots illustrating the gating strategy for singlets (left), total cells (middle), and live cells (right). B) Fluorescence histograms showing lectin binding profiles for Siaα2-3Galβ1-4GlcNAc (MAA-I in FITC, left), Siaα2-3Galβ1-3GalNAc (MAA-II in VioGreen, middle), and Neu5Acα2-6 (SNA in APC, right).

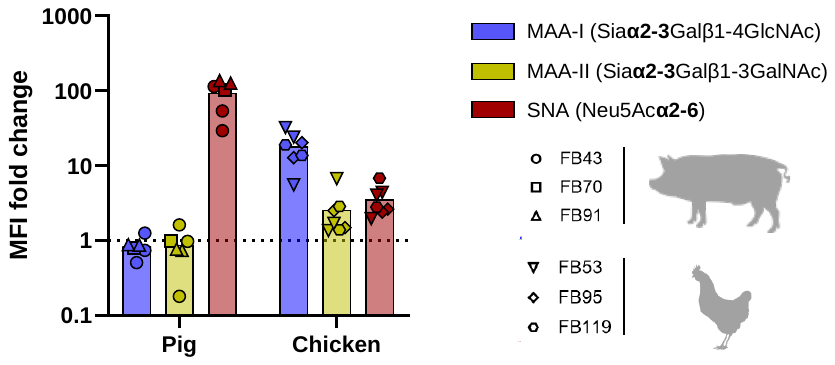

**Supplementary Figure 4. Examining inter-individual variability of sialic acid expression in animals from the same species.** Sialic acid expression using specific lectins against Siaα2-3Galβ1-4GlcNAc [*Maackia amurensis* agglutinin-I (MAA-I, blue), Siaα2-3Galβ1-3GalNAc [*Maackia amurensis* agglutinin-II (MAA-II, yellow)], and Neu5Acα2-6 [*Sambucus nigra agglutinin* (SNA, red)]. Each point represents an individual staining from three different animals per species: FB43 (round), FB70 (square), or FB91 (triangle) for pigs; or FB53 (inverted triangle), FB95 (diamond), or FB119 (hexagon) for chicken.

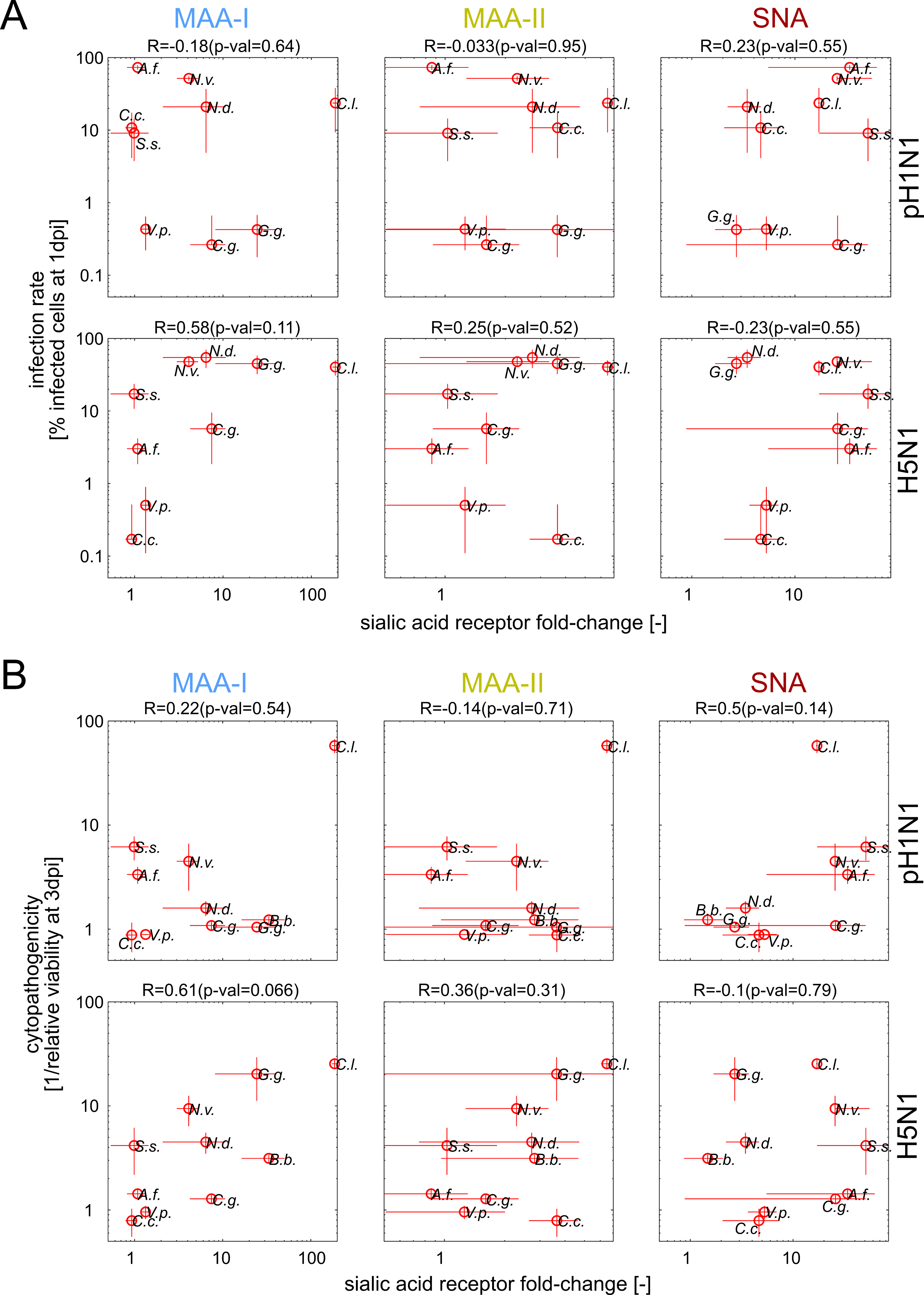

**Supplementary Figure 5. Correlation of Influenza A receptor expression and infection rate at 1 dpi (A) and cytopathogenicity (B) across organoids derived from different animal species.** Sialic acid receptor fold-change (see main Figure 4) plotted against the % infected cells at 1 dpi (**A**), or cytopathogenicity (calculated as 1/relative cell viability at 3 dpi) (**B**) in organoid cultures infected with pH1N1 (top) or H5N1 (bottom). In each comparison, R denotes the Spearman correlation coefficient (with the respective p-value). Each circle denotes an animal (designated with a two-letter code from its Latin name). Error bars denote standard deviation (n = 2-6 infected wells).

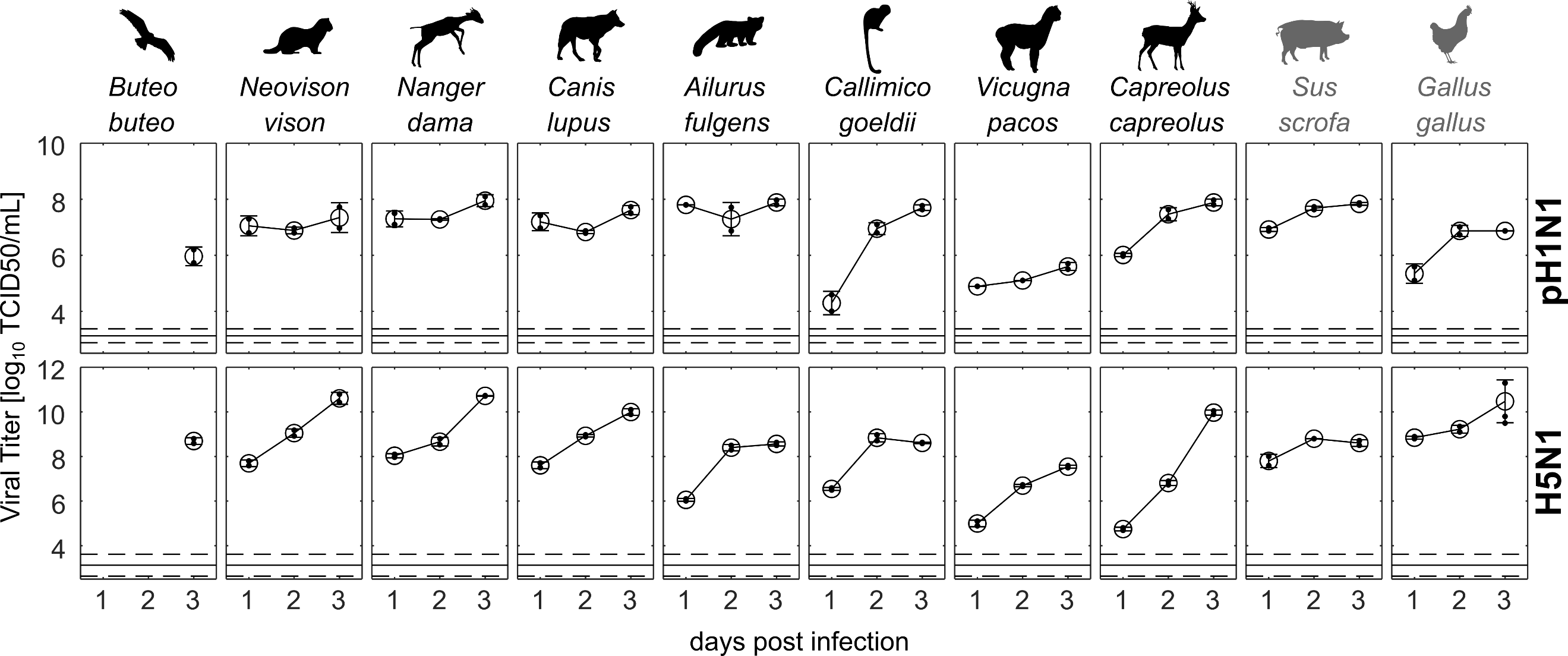

**Supplementary Figure 6. Quantification of infectious viral particles during pH1N1 (top row) and H5N1 (bottom row) infection.** Viral titers in culture supernatants were quantified as described in the methods section. Large circles denote the mean, error bars denote standard deviation (n = 2-3). Small circles denote individual replicates. The horizontal line represents the titer of the inoculum (dashed lines denote its standard deviation).

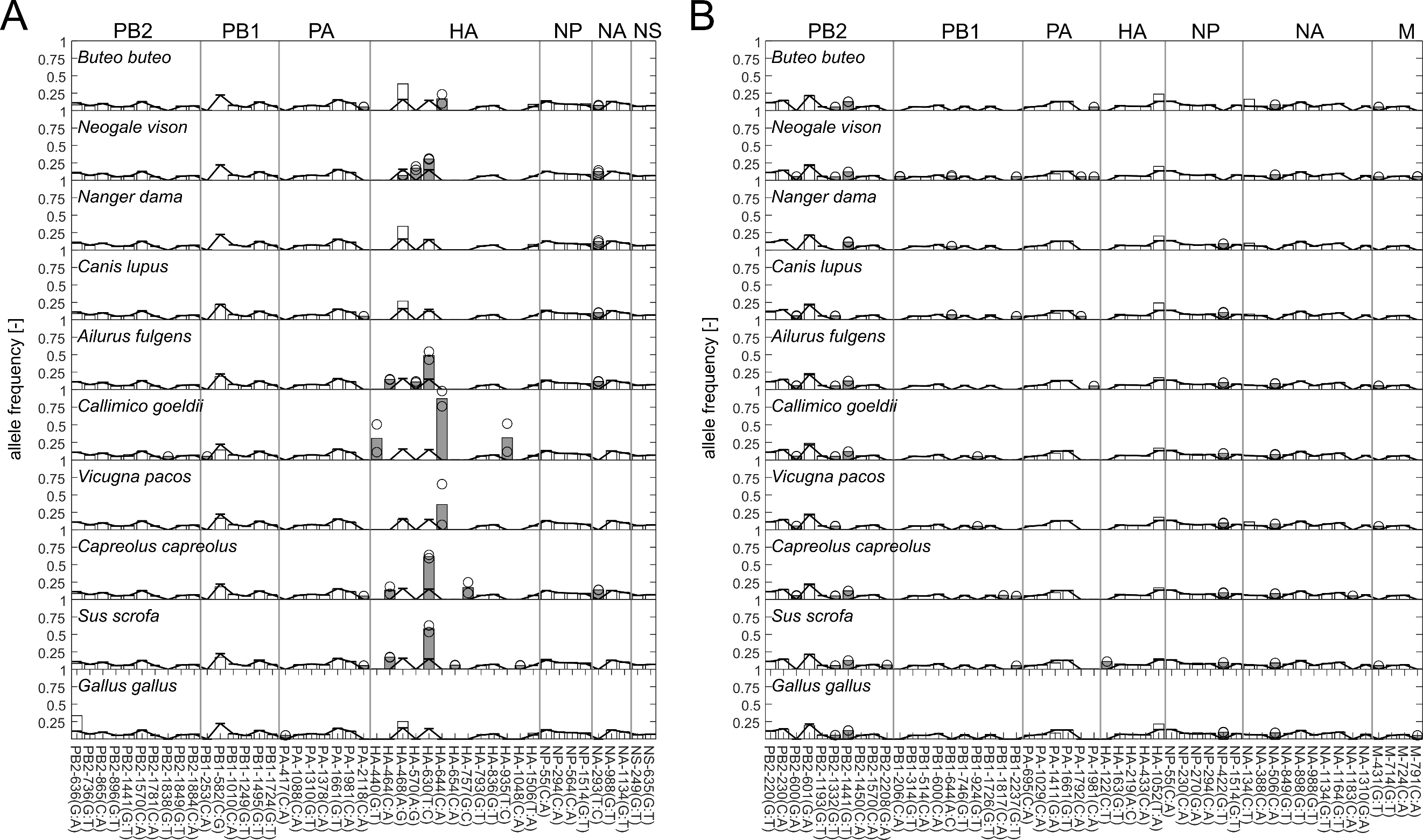

**Supplementary Figure 7. Examining viral adaptation of different organoid cultures infected with pH1N1 (A) or H5N1 (B).** Rows denote animal species, columns denote mutations (sorted based on segment and position). Continuous black line: allele frequency in inoculum. Gray bars: mutations with >2-fold difference in allele frequency in both replicates compared to the inoculum. Black circles: corresponding individual replicates.

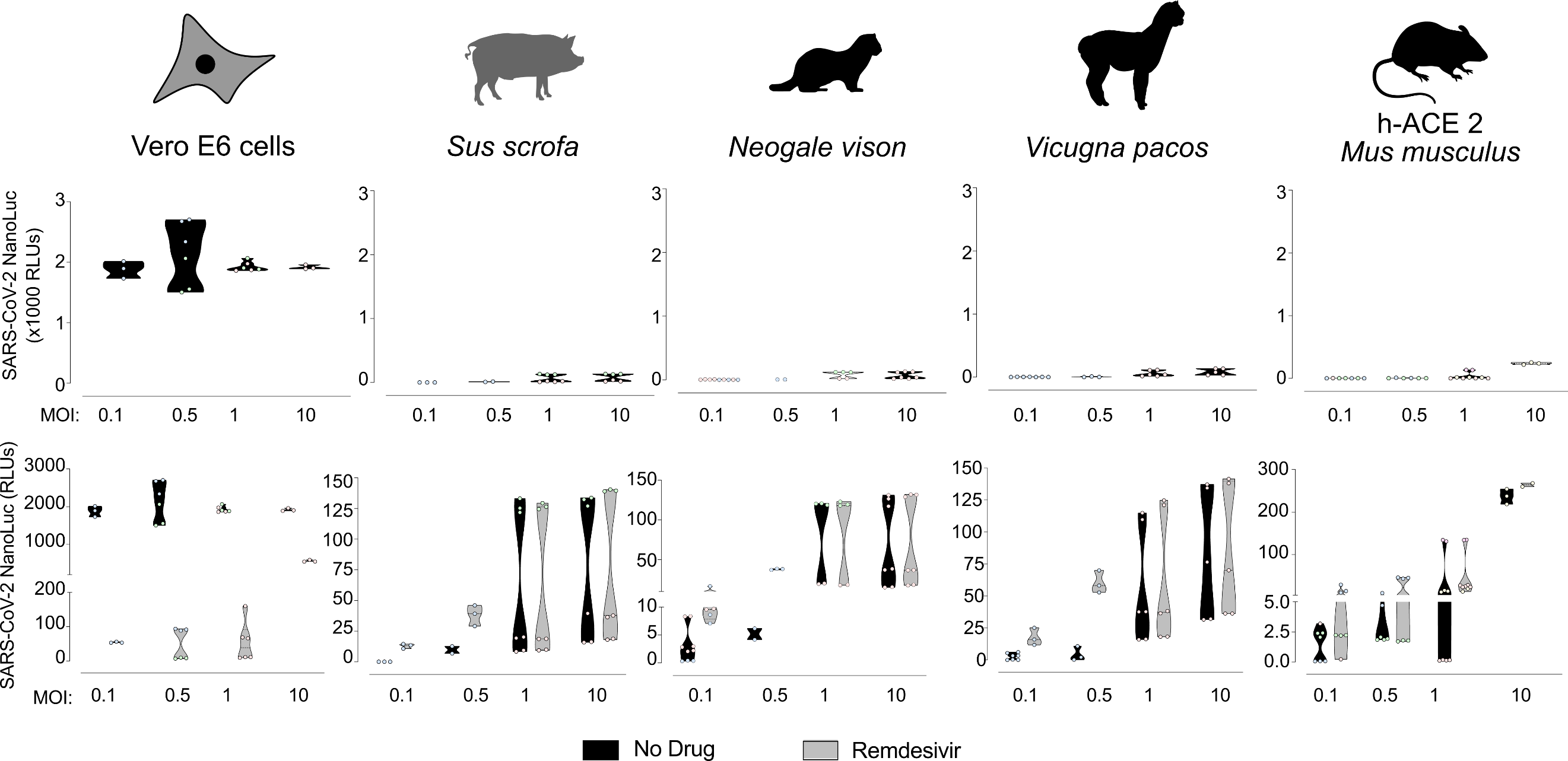

**Supplementary Figure 8. SARS-CoV-2 does not infect animal organoid-derived 2D-monolayers.** Nanoluciferase detected in Vero E6 cells, 2D pig lung organoids, 2D mink trachea organoids, 2D alpaca trachea organoids, 2D hACE2 mice lung organoids 72h post infection at different MOIs using a highly sensitive nanoluciferase SARS-CoV-2 reporter virus. Luminometry values are expressed in relative light units (RLUs). Top row: experiments with virus alone. Bottom row: experiments with virus and Remdesivir (SARS-CoV-2 inhibitor). Data show violin plots with median and quartiles. Cells plotted in light grey plots were treated with Remdesivir at 25µM. Dots represent replicas and colors show individuals tested. Experiments were performed at least 3 times with 2-3 replicas. Background levels of SARS-CoV-2 nanoluciferase inoculum for MOI 0.1 to 10 were 1.14 RLUs (min.) to 212.10 RLUs (max.).
