## Supplementary figures and images for "Airway organoids reveal patterns of Influenza A tropism and adaptation in wildlife species"

### High-resolution version of Figure 5

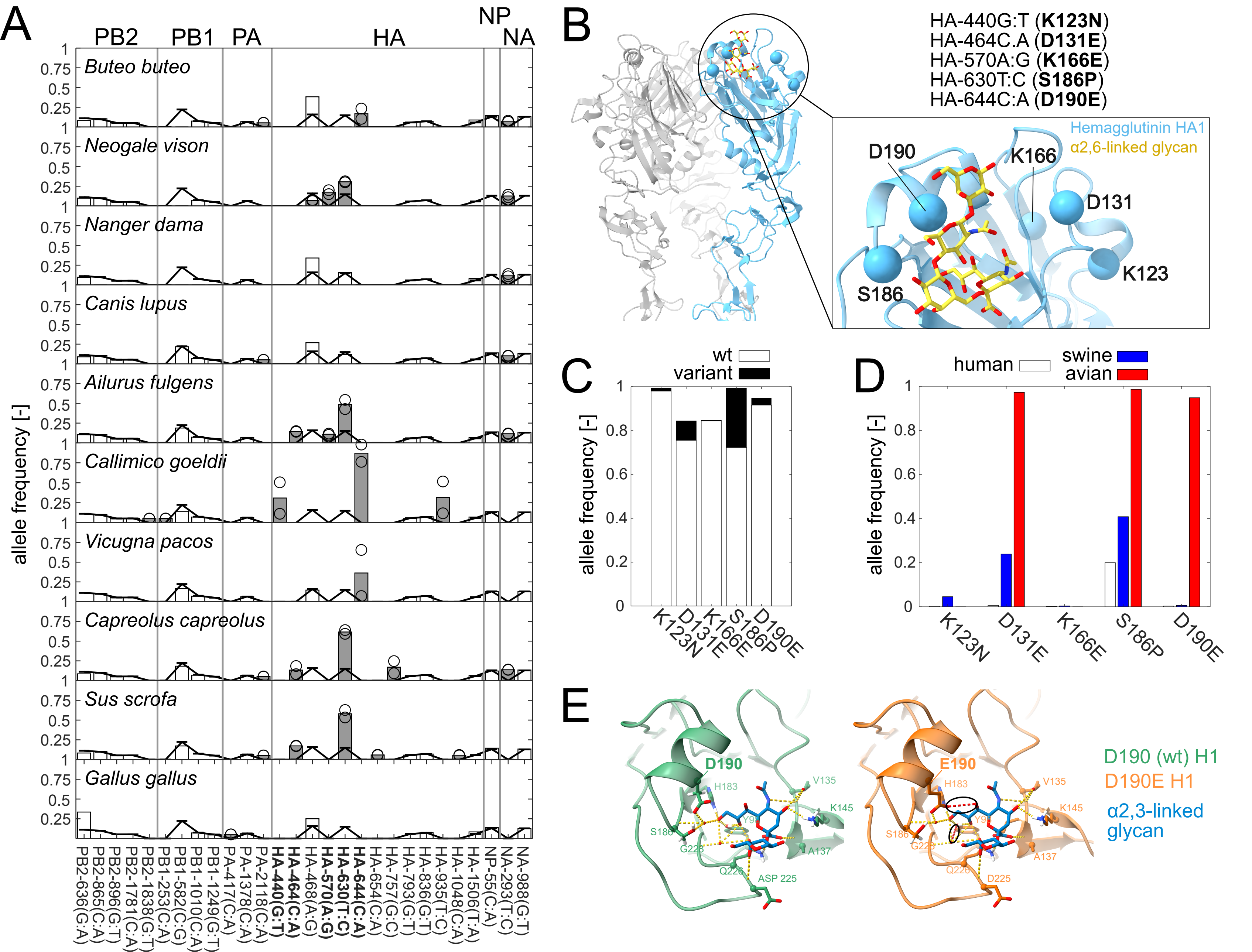
